# High-resolution in situ structures of hantavirus glycoprotein tetramers

**DOI:** 10.1101/2025.06.17.660152

**Authors:** Luqiang Guo, Elizabeth McFadden, Megan M. Slough, E. Taylor Stone, Jacob Berrigan, Eva Mittler, Kiara Hatzakis, Troy Hinkley, Heather S. Kain, Zunlong Ke, Nikole L. Warner, Jesse H. Erasmus, Kartik Chandran, Jason S. McLellan

## Abstract

New World hantaviruses cause severe infections in humans, with case fatality rates approaching 40%. Previous structural studies have advanced our understanding of hantavirus glycoprotein architecture and function, however, the lack of high-resolution in situ structures of the glycoprotein tetramer and its lattice organization has limited mechanistic insights into viral assembly, entry, and antigenicity. Here, we leveraged a virus-like particle (VLP) system to establish a cryo-electron microscopy workflow for lattice-forming viral glycoproteins. This enabled the determination of a 2.35 Å resolution structure of the membrane-embedded Andes virus (ANDV) glycoprotein tetramer, as well as structures of dimers of tetramers and a complex with antibody ADI-65534. These structures reveal previously uncharacterized features of glycoprotein organization, stability, and pH-sensing. Immunization of mice with self-amplifying replicon RNA (repRNA) encoding ANDV-VLPs elicited high levels of glycoprotein-binding antibodies but equivalent titers of neutralizing antibodies compared to repRNA-encoded native ANDV glycoprotein complex. Collectively, these findings advance our understanding of hantavirus glycoprotein assemblies and their function, laying a foundation for structure-based vaccine design efforts.

## INTRODUCTION

Andes virus (ANDV), a rodent-borne New World hantavirus, causes hantavirus cardiopulmonary syndrome (HCPS) in humans, with case fatality rates reaching 40% (Vial, Ferrés et al. 2023). Hantaviruses are classified into Old World hantaviruses (OWHs), which cause hemorrhagic fever with renal syndrome (HFRS), and New World hantaviruses (NWHs), which cause the more frequently fatal HCPS. Human infection occurs primarily through inhalation of aerosolized rodent saliva and excreta, although human-to-human transmission through close contact has been documented for ANDV (Padula, Edelstein et al. 1998, Martinez-Valdebenito, Calvo et al. 2014, Martínez, Di Paola et al. 2020). Despite the public health threat posed by ANDV, there are no approved vaccines or therapeutics.

Like other viruses of the family *Hantaviridae*, ANDV is an enveloped virus enclosing a tri-segmented, negative-strand RNA genome (Cifuentes-Muñoz, Salazar-Quiroz et al. 2014, Mittler, Dieterle et al. 2019). The medium (M) segment encodes a glycoprotein precursor (GPC), which is co-translationally cleaved by signal peptidase in the endoplasmic reticulum (ER) into two glycoproteins, Gn and Gc (Löber, Anheier et al. 2001). These proteins form a stable heterodimer that oligomerizes into tetramers that form a curved lattice structure critical for virion assembly and budding (Huiskonen, Hepojoki et al. 2010, Serris, Stass et al. 2020). Hantavirus assembly predominantly occurs at the Golgi apparatus or plasma membrane (Goldsmith, Elliott et al. 1995, Xu, Yang et al. 2007, Barker, daSilva et al. 2023). Following egress, ANDV binds to its primary receptor, PCDH1, for entry (Jangra, Herbert et al. 2018). Following endocytosis, the acidic endosomal environment triggers conformational changes in the glycoprotein complex (Acuña, Bignon et al. 2015). This leads to release of the class II viral fusion protein, Gc, from the Gn–Gc complex (Rissanen, Stass et al. 2017), allowing Gc to mediate fusion of the viral and endosomal membranes as it transitions from the metastable prefusion conformation to the stable postfusion conformation (Cifuentes-Muñoz, Salazar-Quiroz et al. 2014, Mittler, Dieterle et al. 2019).

Gn and Gc are the only surface-exposed glycoproteins on hantavirus virions and serve as the primary targets of the neutralizing antibody response (Dantas, Okuno et al. 1986, Arikawa, Schmaljohn et al. 1989). Antibodies that cross-neutralize multiple hantaviruses have been reported (Chu, Jennings et al. 1995, Lundkvist, Hukic et al. 1997), and survivors of hantavirus infection often retain long-lasting neutralizing antibody titers (Valdivieso, Vial et al. 2006). Moreover, neutralizing antibody titers are a strong correlate of protection for both HFRS and HCPS patients (Bharadwaj, Nofchissey et al. 2000, MacNeil, Comer et al. 2010, Pettersson, Thunberg et al. 2014). These observations have spurred extensive vaccine development efforts aimed at eliciting robust, long-lasting, and broadly neutralizing immune responses directed toward the Gn and Gc antigens (Afzal, Ali et al. 2023, Sehgal, Mehta et al. 2023). Notably, recombinant vesicular stomatitis viruses (rVSVs) expressing ANDV (Brown, Safronetz et al. 2011) or Sin Nombre virus (SNV) glycoproteins (Warner, Stein et al. 2019) have shown efficacy in animal models. Additionally, an ANDV M segment-based DNA vaccine (Custer, Thompson et al. 2003) has demonstrated protective effects. These approaches induced cross-neutralizing antibodies and protected hamsters and rhesus macaques against lethal challenge with ANDV and SNV, respectively.

In parallel, high-throughput isolation and characterization of monoclonal antibodies have advanced our understanding of the humoral immune response to hantavirus infection (Garrido, Prescott et al. 2018, Engdahl, Kuzmina et al. 2021, Mittler, Wec et al. 2022). Structural studies using X-ray crystallography and cryo-electron tomography (cryo-ET) have provided insights into the molecular mechanisms of antibody-mediated neutralization (Rissanen, Stass et al. 2020, Rissanen, Krumm et al. 2021, Mittler, Wec et al. 2022, Engdahl, Binshtein et al. 2023, Stass, Engdahl et al. 2023) . However, high-resolution structural information of antibodies in complex with Gn–Gc in their tetrameric or lattice-associated forms remains limited.

Structural studies of authentic ANDV virions have been constrained by Biosafety Level 3 requirements (Parvate, Williams et al. 2019). Nevertheless, significant insights have been obtained from studies of apathogenic hantaviruses, like Tula virus (TULV). In addition, investigations of Old World hantaviruses, including Puumala virus (PUUV) and Hantaan virus (HTNV), have elucidated the organization of the glycoprotein lattices and the conformational arrangement of Gn and Gc (Martin, Lindsey-Regnery et al. 1985, Huiskonen, Hepojoki et al. 2010, Guardado-Calvo, Bignon et al. 2016, Li, Rissanen et al. 2016, Willensky, Bar-Rogovsky et al. 2016, Rissanen, Stass et al. 2017). A model of the ANDV glycoprotein tetramer and lattice was subsequently generated by docking crystal structures of the ANDV Gn base tetramer and a single-chain construct of the Gn head and Gc ectodomain heterodimer in its prefusion conformation into the cryo-ET map of TULV (Serris, Stass et al. 2020). In this model, Gn resides centrally, mediating tetramerization, whereas Gc is positioned peripherally along the edge of each tetramer, mediating tetramer–tetramer interactions. However, differences in tetramer architecture between OWHs and NWHs complicate direct extrapolation to ANDV (Battisti, Chu et al. 2011), and the absence of high-resolution structures of the tetramers in their native membrane environment continues to impede a comprehensive molecular understanding of ANDV architecture, function, and antibody-mediated inhibition.

Here, we demonstrate that the addition of an eVLP tag (Hoffmann, Yang et al. 2023) to the ANDV GPC substantially enhances the production of ANDV-VLPs. Purification of these VLPs enabled single-particle cryo-EM studies, allowing us to determine the structure of individual ANDV Gn–Gc tetramers to 2.35 Å resolution, as well as dimers of ANDV Gn–Gc tetramers in three related flexing conformations to 3.2 Å, 3.4 Å, and 3.4 Å resolution. Furthermore, we resolved the structure of the antigen-binding fragment (Fab) of ADI-65534 (Mittler, Serris et al. 2023), an engineered pan-hantavirus antibody, in complex with ANDV tetramers and dimers of tetramers, unexpectedly demonstrating that the full-length IgG is unable to cross-link neighboring tetramers. These structures reveal the molecular basis of Gn–Gc tetramer organization, lattice formation, acid-induced membrane fusion, and antibody-mediated neutralization. Additionally, immunogenicity studies of ANDV-VLPs as a self-amplifying replicon RNA (repRNA) vaccine candidate revealed improved binding—but equivalent neutralizing—antibody titers, suggesting a need to further characterize determinants of repRNA-encoded ANDV glycoprotein immunogenicity.

## RESULTS

### Characterization of ANDV rVSVs and VLPs

We initially attempted to use the established rVSV-ANDV system (Kleinfelter, Jangra et al. 2015) to structurally characterize the ANDV glycoprotein lattice in a membrane context (**Fig. 1A**). Cryo-electron microscopy (cryo-EM) screening of untreated rVSV-ANDV particles revealed characteristic bullet-shaped virions with glycoprotein densities restricted to the curved tips (**Fig. 1B–C, S1A**), contrasting with the more uniform distribution observed for VSV G proteins on authentic virions (Zhou, Si et al. 2022, Xin, Kurien et al. 2025). To assess whether this polarized localization is unique to ANDV glycoproteins, we examined rVSV particles pseudotyped with the SARS-CoV-2 spike protein. Unlike ANDV Gn–Gc, the SARS-CoV-2 spike protein exhibited an even distribution across the viral envelope (**Fig. 1B–C, S1B**), albeit in a predominantly postfusion conformation, suggesting that the tip-localization observed for ANDV glycoproteins is not an inherent feature of the VSV backbone. Given the phylogenetic divergence between ANDV and SARS-CoV-2, we next evaluated the glycoprotein distribution in rVSV particles bearing glycoproteins from Crimean–Congo hemorrhagic fever virus (CCHFV), which is a member of the family *Nairoviridae* and, like ANDV, is a bunyavirus (Simmonds, Adriaenssens et al. 2024). Similar to ANDV, CCHFV glycoproteins predominantly localized to the tips of the rVSV virions (**Fig. S1C, S1D**), indicating that this spatial preference may represent a conserved feature among bunyaviruses.

**Fig. 1.**
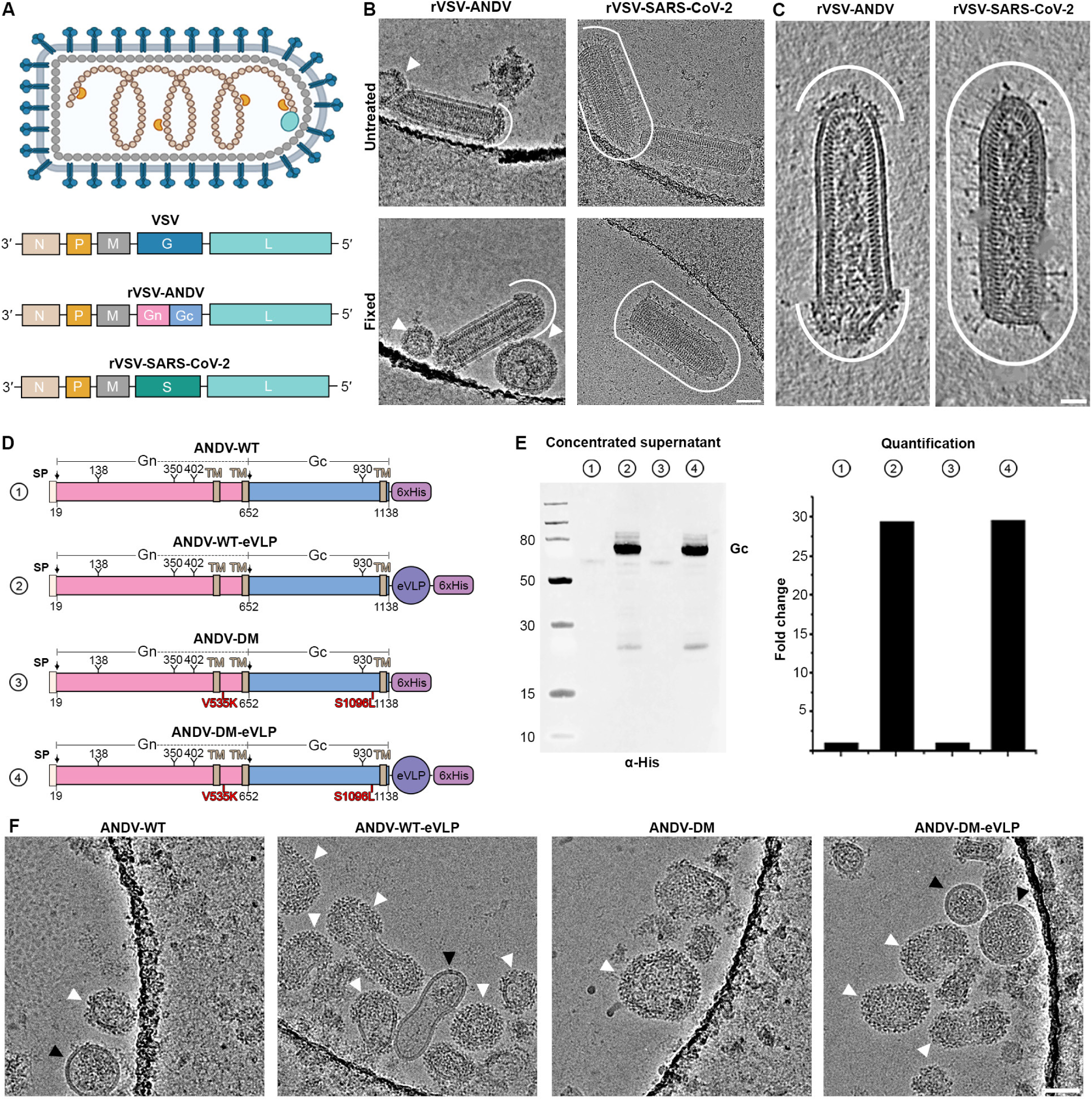
The eVLP system substantially improves the yield of ANDV VLPs for cryo-EM imaging. (**A**) Diagram of the VSV virion and its genome organization. rVSV-ANDV and rVSV-SARS-CoV-2 were generated by replacing the gene encoding VSV G with the genes encoding ANDV and SARS-CoV-2 glycoproteins. (**B**) Cryo-EM images of untreated and glutaraldehyde-fixed rVSV-ANDV and rVSV-SARS-CoV-2 samples. White curved lines indicate the distribution of glycoproteins on the virion surface. VLPs displaying ANDV glycoproteins are denoted by white arrowheads. Scale bar: 50 nm. (**C**)16-nm-thick tomography slices of untreated rVSV-ANDV and rVSV-SARS-CoV-2 samples. White curved lines indicate the distribution of glycoproteins on the virion surface. Scale bar: 25 nm. (**D**) Schematics of ANDV Gn-Gc expression constructs. SP: signal peptide; TM: transmembrane domain; Y: N-linked glycosylation site; eVLP: EPM and EABR sequences. (**E**) Western blot analysis of concentrated supernatants from cells transfected with the constructs in (D). Bands corresponding to Gc (*left*) were quantified and graphed as the fold-change in expression compared to the ANDV-WT construct (*right*). (**F**) Cryo-EM images of purified ANDV-eVLPs. White arrowheads denote eVLPs displaying ANDV glycoproteins. Black arrowheads denote VLPs lacking glycoprotein lattices. Scale bar: 50 nm.

Due to the low abundance of rVSV-ANDV virions and the dense glycoprotein coverage observed on the surface of the VLPs, we elected to pursue high-resolution structural studies of the ANDV VLPs. Previous studies have demonstrated that expression of the full M segment-encoded GPC is sufficient to drive self-assembly of ANDV glycoproteins into VLPs (Acuña, Cifuentes-Muñoz et al. 2014). Following this protocol, we attempted to generate ANDV-VLPs from small-scale transfections; however, these initial efforts were unsuccessful due to insufficient particle yields (**Fig. 1D–E**). To improve expression and budding efficiency, we incorporated two substitutions, V535K and S1096L, which have been previously shown to enhance the plasma membrane localization of HTNV glycoproteins (Slough, Chandran et al. 2019). We also added a C-terminal eVLP tag, which facilitates budding of VLPs from the cell membrane by recruiting components of the ESCRT pathway (Hoffmann, Yang et al. 2023) (**Fig. 1D**). Western blot analysis of concentrated supernatants revealed that the double modification (ANDV-DM) had a minimal impact on glycoprotein yield, whereas the eVLP tag (ANDV-WT-eVLP) enhanced the yield approximately 30-fold relative to wild-type ANDV (ANDV-WT) (**Fig. 1E**). The combination of both modifications (ANDV-DM-eVLP) did not substantially improve expression over the ANDV-WT-eVLP construct (**Fig. 1E**). Cryo-EM screening of purified preparations from each construct further corroborated these findings: ANDV-WT-eVLP and ANDV-DM-eVLP yielded abundant VLPs in nearly every micrograph, whereas ANDV-WT and ANDV-DM samples contained only sparse VLPs (**Fig. 1F**). As cryo-EM grids for ANDV-DM-eVLPs appeared qualitatively better than the ANDV-WT-eVLP grids, further structural studies were performed on ANDV-DM-eVLP samples.

### Overview of the in situ ANDV glycoprotein tetramer architecture

With the substantially improved VLP yield, and in light of recent advances in single-particle structure determination of proteins embedded in membranes (Ke, Oton et al. 2020, Tao, Zhao et al. 2023, Calcraft, Stanke-Scheffler et al. 2024, Karimi, Coupland et al. 2024), we pursued single-particle analysis (SPA) cryo-EM studies of the ANDV glycoprotein lattice on ANDV-DM-eVLPs. However, the pleomorphism of the particles and lattice, combined with overlapping projections of glycoprotein tetramers, posed challenges for conventional SPA processing. Following systematic optimization—including refinement of particle-picking strategies, adjustment of extraction box sizes, and particle recentering—we obtained high-quality 2D class averages displaying clear secondary structure features, including the transmembrane regions (**Fig. 2A, S2**). Subsequent rounds of 3D classification and refinement yielded a 2.35 Å resolution reconstruction of the ANDV glycoprotein tetramer, refined with C4 symmetry, from a dataset comprising 253,514 particles extracted from 17,681 movies (**Fig. 2B, S2 and S3A–B**).

**Fig. 2.**
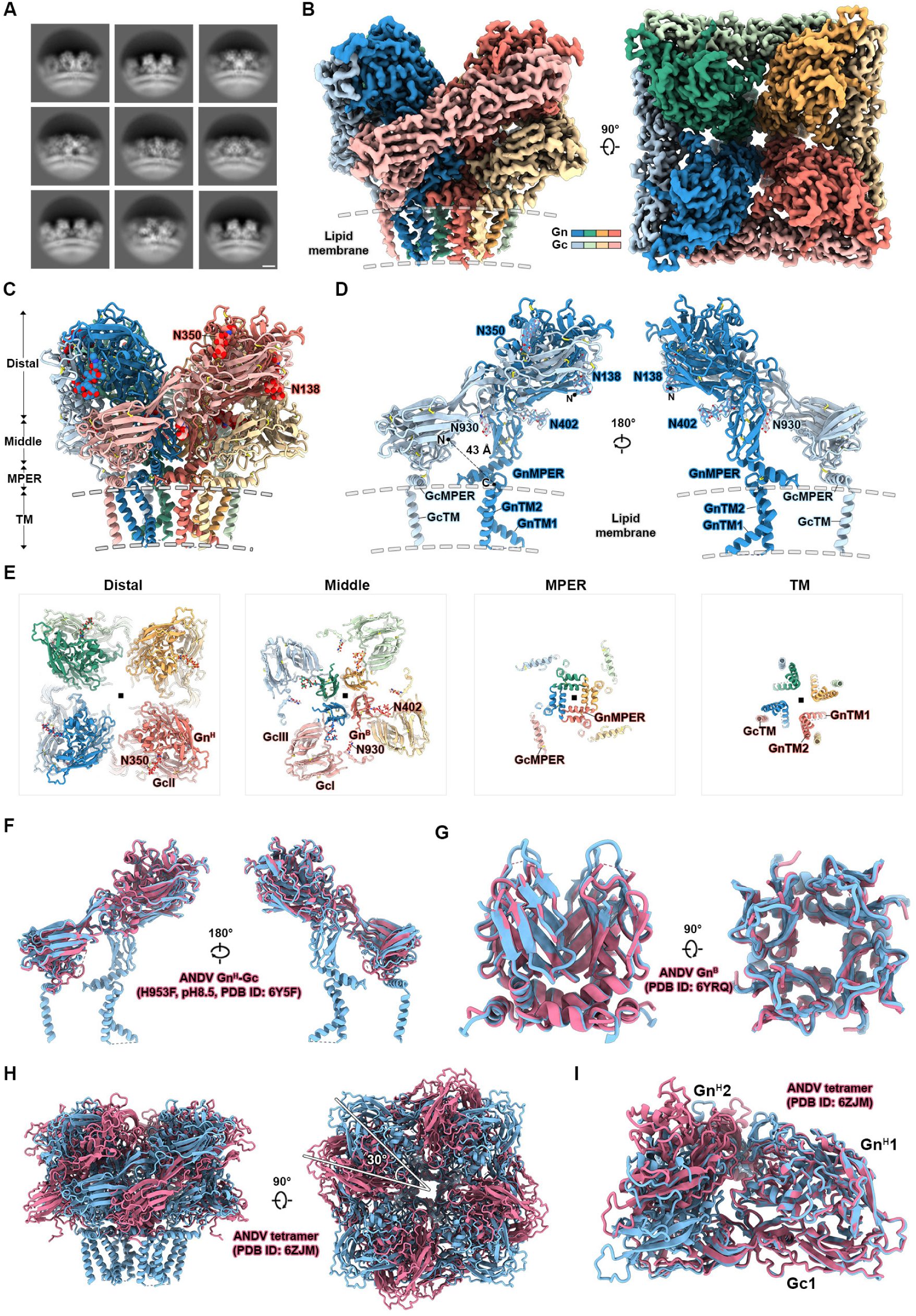
Architecture of the ANDV Gn–Gc tetramer in a membrane. (**A**) Cryo-EM 2D class averages of ANDV glycoproteins on the surface of eVLPs. Scale bar: 50 Å. (**B**) Cryo-EM map processed by DeepEMhancer of the membrane-bound ANDV Gn–Gc tetramer. Gn–Gc heterodimers are two shades of the same color. (**C**) Structural model of the ANDV Gn–Gc tetramer, shown as ribbons and colored as in (B). N-linked glycan atoms are shown as spheres. Nitrogen atoms are colored dark blue, oxygen atoms are colored red, and sulfur atoms are colored yellow. MPER: membrane-proximal external region; TM: transmembrane domain. (**D**) Model of a Gn–Gc heterodimer, shown as ribbons and colored as in (B). N-linked glycans are shown as sticks with their corresponding cryo-EM map densities displayed as partially transparent surfaces. The black dashed line indicates the distance between the C-terminal and N-terminal residues of Gn and Gc, respectively. (**E**) Slices through a top-down view of the ANDV Gn–Gc tetramer shown as ribbons and colored as in (B). Domains and features of one protomer are labeled in each panel. (**F**) A Gn–Gc heterodimer from the cryo-EM structure (blue) aligned to a previously determined crystal structure of an ANDV Gn–Gc fusion construct (PDB ID: 6Y5F, pink). (**G**) Alignment of the GnB domain from the cryo-EM structure (blue) to a previously determined crystal structure of the ANDV GnB domain (PDB ID: 6YRQ). (**H**) The ANDV Gn–Gc tetramer from the in situ cryo-EM structure (blue) aligned to a model previously obtained by fitting the crystal structures of ANDV Gn–Gc and ANDV GnB into an 11 Å resolution subtomogram average of the Tula virus Gn– Gc tetramer (pink, PDB ID: 6ZJM). (**I**) GnH1–Gc1 and GnH2 from the in situ cryo-EM structure (blue) and the published tetramer model (pink) aligned via GnH1.

The high-resolution cryo-EM reconstruction revealed well-defined sidechain densities for most amino acid residues, along with densities corresponding to glycan chains at all four predicted N-linked glycosylation sites. These features facilitated unambiguous model building (**Fig. 2C, S3C–F**), resulting in a final model encompassing the complete ectodomains of both Gn and Gc and their transmembrane domains (**Fig. 2C–D**). Residues 513–627 (Gn endodomain), 651–652 (N-terminus of Gc), and 1128–1138 (Gc endodomain) were not resolved in the map, likely due to their intrinsic flexibility and the absence of ribonucleoprotein (RNP) complexes that may stabilize these regions (Hepojoki, Strandin et al. 2010, Wang, Alminaite et al. 2010). The final model also includes 452 water molecules, several of which participate in hydrogen-bonding networks that appear to stabilize inter-subunit interactions within the tetramer. Notably, during model building, we identified several unassigned densities that may correspond to lipids or post-translational modifications (**Fig. S4**).

The membrane-distal region of the tetramer is primarily composed of Gc domain II (GcII) and the Gn ‘head’ domain (Gn^H^), including N-linked glycans at Asn138 and Asn350 on Gn^H^ (**Fig. 2E**). The middle region is composed of Gc domain I (GcI), parts of GcII, most of Gc domain III (GcIII), and the majority of the Gn ‘base’ domain (Gn^B^). The other two N-linked glycans, at Asn402 on Gn and Asn930 on Gc, are also in this region. The glycan chains are not on the surface of the tetramer, but are internal, and may play a structural role, as noted by Serris et al (Serris, Stass et al. 2020). The membrane-proximal external region (MPER) contains two amphipathic helices in a helix-turn-helix motif from each Gn subunit, forming a square scaffold that supports the tetramer on the membrane. This region also contains an amphipathic helix from Gc, although this helix does not interact with the Gn MPER helices. The transmembrane (TM) region contains two helices from Gn and one helix from Gc. The first Gn TM helix (GnTM1) is kinked approximately 90° midway along its length. The Gc TM helices (GcTM) are located on the exterior of this arrangement, with a TM helix from Gc interacting with the Gn TM helices of an adjacent protomer. The molecular interactions in each of these regions are described in more detail in the following Results section.

Structural comparison between the ANDV Gn–Gc heterodimer from the high-resolution tetramer model and the previously determined crystal structure of the prefusion Gn^H^–Gc complex (PDB ID: 6Y5F) revealed minimal conformational differences in the overall architecture of both Gn^H^ and Gc, with a root mean square deviation (RMSD) of 1.1 Å for 769 Cα atoms when aligned on Gc domain II (**Fig. 2F**). Superposition of the Gn^B^ domains (PDB ID: 6YRQ) also revealed close agreement, with an RMSD of 1.0 Å for 396 Cα atoms (**Fig. 2G**). The published ANDV tetramer model (PDB ID: 6ZJM), constructed by fitting the Gn^H^–Gc and Gn^B^ crystal structures into an 11 Å subtomogram average of the TULV glycoprotein tetramer, is missing multiple regions present in our in situ tetramer structure, including the N terminal loop of Gc (residues 653–661), the stem loop within domain III of Gc (residues 1063–1076), both Gn transmembrane helices (GnTM1: residues 483–512 and GnTM2: residues 628–650), as well as the GcMPER and transmembrane domain (GcTM; residues 1093–1127). Structural alignment of the two tetramer models based on the Gn^B^ domains revealed an approximate 30° rotation of the Gn^H^–Gc protomers (**Fig. 2H**), resulting in an incorrect spatial positioning of the Gn^H^–Gc protomers relative to each other as well as to Gn^B^ in the published model. This is best visualized in **Supplemental Movie 1**. This misalignment can be alternatively viewed by superposition of a Gn^H^1–Gc1 protomer on Gn^H^1 and observing the relative position of an adjacent Gn^H^2, which demonstrates the displacement of Gn^H^2 from the Gn^H^1–Gc1 protomer (**Fig. 2I**). Critically, the incorrect orientation in the previously published model eliminates two key inter-subunit interfaces that mediate Gn2–Gc1 interactions, described in more detail in the subsequent Results section. The observed misplacement likely reflects genuine structural variability among hantavirus tetramers, as previously suggested by comparisons between HTNV and TULV based on low-resolution EM maps (Battisti, Chu et al. 2011).

### Molecular interfaces stabilizing the ANDV glycoprotein tetramer

The in situ ANDV glycoprotein tetramer structure enables residue-level interrogation of inter- and intra-protomer interactions, offering molecular insights into tetramer assembly. Analysis of inter-subunit contacts reveals three principal interactions that stabilize the assembly: Gn1–Gc1, Gn1–Gn2, and Gn2–Gc1 (**Fig. 3**).

**Fig. 3.**
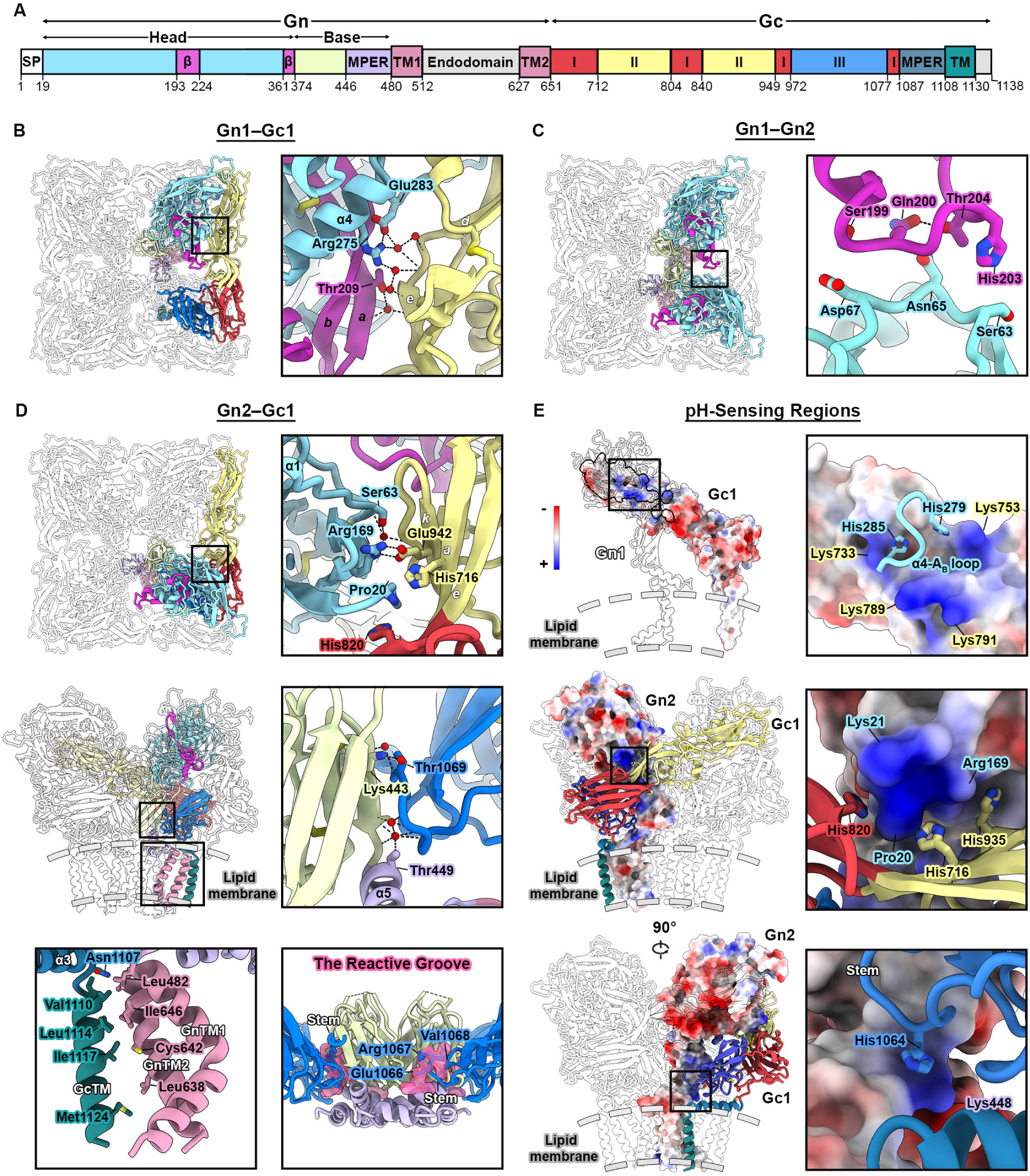
Novel interfaces that stabilize the ANDV glycoprotein tetramer. (**A**) Schematic of the ANDV M segment, colored by domain. (**B**) Top-down view of the in situ ANDV Gn–Gc tetramer structure shown as ribbons (left). Portions of Gn1 and Gc1 are colored by domain, as in (A). Zoomed-in view of the Gn1–Gc1 interface (right). (**C**) The Gn1–Gn2 interface, depicted as in (B). (**D**) The Gn2–Gc1 interfaces, depicted as in (B). The reactive groove (bottom right) is shown as transparent pink surfaces. (**E**) Histidine-containing interfaces that are likely involved in destabilization of the Gn–Gc tetramer upon acidification in endosomes. The Gn1–Gc1 interface (top) with Gn1 shown as transparent ribbons and Gc1 shown as a molecular surface colored by electrostatic potential. The zoomed-in panel shows Gn histidine sidechains as sticks. The membrane-distal region of the Gn2–Gc1 interface (middle) with Gn2 shown as a molecular surface colored by electrostatic surface potential. The membrane-proximal Gn2–Gc1 interface. In all panels, nitrogen atoms are colored blue, oxygen atoms are red, sulfur atoms are yellow, water molecules are shown as red spheres, and hydrogen bonds are shown as black dashed lines.

### Gn1***–***Gc1 interface

The Gn1–Gc1 interface is primarily mediated by contacts between Gn^H^1 and domain II of Gc1 (Gn^H^1–Gc1II), along with two key N-linked glycans at Asn138 and Asn350 on Gn^H^1, which together form a clamp-like structure that secures Gc1 in the appropriate orientation (**Fig. 2C-D**), consistent with prior findings (Serris, Stass et al. 2020). This interface buries 1,884 Å^2^ of solvent-accessible surface area when glycans are included and 1,437 Å^2^ without. Notably, the high-resolution map reveals a hydrogen bond network at the Gn^H^1–Gc1II interface, involving side chains of Thr209, Arg275, and Glu283 of Gn^H^1, backbone atoms of β-strand *e* and its upstream loop in Gc1, and several ordered water molecules, which collectively contribute to interface stabilization **(Fig. 3A-B**).

### Gn1***–***Gn2 interface

The Gn1–Gn2 interaction comprises two major interfaces, Gn^H^1–Gn^H^2 (**Fig. 3C**) and Gn^B^1–Gn^B^2 (**Fig. 2G**). The Gn^H^1 and Gn^H^2 interface is formed by the loop upstream of β-strand *a* of Gn^H^1 and the loop downstream of α1 of Gn^H^2, which buries a total surface area of 176 Å^2^ and mainly involves mainchain-mainchain and mainchain-sidechain interactions (**Fig. 3C)**. This interface was distorted in the previous ANDV tetramer model (PDB ID: 6ZJM). The Gn^B^1–Gn^B^2 interface is essentially the same as that described previously for the Gn^B^ crystal structure, here burying a total solvent-accessible surface area of 963 Å^2^.

### Gn2***–***Gc1 interface

The Gn2–Gc1 interactions are previously uncharacterized and can be subdivided into three spatially distinct interfaces: the Gn^H^2–Gc1 interface, the Gn^B^2–Gc1 interface, and the transmembrane interaction between Gn2 and Gc1 (Gn2TMs–Gc1TM) (**Fig. 3D**). The Gn^H^2–Gc1 interface buries 406 Å^2^ of surface area and involves contacts between Gn^H^2 and domain II of Gc1 (**Fig. 3D, top panels**). This interface may be important for tetramer dissociation under acidic conditions (**Fig. 3E, middle panels**), as the sidechains of His716, His820, and His935 on Gc1 are adjacent to a highly positively charged surface on Gn^H^2 created by Lys21, Arg169, and the amino-terminal residue Pro20. The Gn^B^2–Gc1 interface buries 455 Å^2^ and is supported by two water-mediated hydrogen bond networks involving Gn^B^2 and the N-terminal stem peptide of Gc1 domain III (Gc1III) (**Fig. 3D, middle panels**). Notably, a reactive groove identified in the previously solved Gn^B^ crystal structure (PDB ID: 6YRQ) was hypothesized to accommodate the N-terminal segment of the Gc stem peptide, which remained unresolved in the Gn^H^–Gc crystal structure due to the absence of Gn^B^ (Serris, Stass et al. 2020). Our in situ structure of the ANDV glycoprotein tetramer provides partial confirmation of this hypothesis: residues 1066–1068 of the Gc stem are observed resting atop this groove (**Fig. 3D, lower-right panel**). Finally, the Gn2TMs–Gc1TM interface is primarily stabilized by hydrophobic interactions between residues within Gn2TM2 and Gc1TM1, burying 226 Å^2^ of solvent-accessible surface area (**Fig. 3D, lower-left panel**). Together, these three Gn2–Gc1 interfaces contribute significantly to the architectural integrity of the tetramer and may play regulatory roles during viral fusion activation.

### pH-sensing interfaces

Given the well-documented role of histidine residues as pH sensors that trigger conformational changes in viral proteins (Carneiro, Stauffer et al. 2003, Bressanelli, Stiasny et al. 2004, Stevens, Corper et al. 2004, Kampmann, Mueller et al. 2006, Peukes, Xiong et al. 2020) and acid-induced membrane fusion for ANDV glycoproteins (Acuña, Bignon et al. 2015), we examined the ANDV tetramer model for histidines in proximity to positively charged residues or other histidines. Strikingly, several such residues were identified within key Gn–Gc interfaces (**Fig. 3E**). In the Gn^H^1–Gc1 interface, although no Gc histidines face a distinct positively charged Gn patch, two Gn histidines—His279 and His285—reside in the α4-A_B_ loop, form intramolecular contacts, and are positioned within a triangular pocket of four Gc lysine residues (Lys733, Lys753, Lys789, Lys791) (**Fig. 3E, upper panels**). At the Gn^H^2–Gc1 interface, three Gc histidines (His716, His820, His935) face the most positively charged region on Gn^H^2, comprising Lys21, Arg169, and the N-terminal Pro20 (**Fig.3E, middle panels**). Additionally, at the Gn^B^2–Gc1 interface, His1064 from the Gc stem peptide is positioned near Lys448 on Gn^B^2 (**Fig. 3E, lower panels**). Protonation of these histidines under acidic conditions would generate electrostatic repulsion, destabilizing their respective interfaces and promoting Gn–Gc dissociation. This cascade of histidine-triggered rearrangements facilitates the rearrangement of the tetramer and refolding of Gc into the stable postfusion trimer, ultimately driving membrane fusion (Harrison 2015, Guardado-Calvo and Rey 2021, White, Ward et al. 2023).

### Molecular basis of lateral interactions between neighboring tetramers

To elucidate the molecular basis of ANDV glycoprotein tetramer lattice formation, we re-extracted particles using an expanded box size and obtained 3D reconstructions for ANDV dimers of tetramers in three closely related yet distinct conformations—designated conformations I, II, and III—at resolutions of 3.2 Å, 3.4 Å, and 3.4 Å, respectively (**Fig. 4, S5, S6**). The three conformations are related by a hinging motion at the tetramer-tetramer interface, best visualized in **Supplementary Movie 2**. Rigid-body fitting of the 2.35 Å tetramer structure into each of the dimer maps yielded excellent refinement statistics (**Table S1**), as expected given that the single tetramer model was obtained using the same particles from two subunits of the dimer of tetramers. Analysis of conformation I revealed that the two tetramers have a lateral displacement of 22 Å and have their 4-fold axes tilted by ∼22° with respect to each other (**Fig. 4A-C**). Structural superposition of all three dimer-of-tetramers conformations revealed progressive flexing between tetramers, with tilt angles of approximately 25° and 27° for conformations II and III, respectively (**Fig. 4C**), suggesting inherent conformational plasticity within the lattice that may be required to accommodate the pleiomorphic virions. Furthermore, if the tilted tetramers represent the lowest-energy conformation of the dimer compared to untilted tetramers, then the adoption of these tilted conformations is consistent with the role for the glycoproteins in inducing membrane curvature and promoting virion or VLP self-assembly, as previously proposed (Huiskonen, Hepojoki et al. 2010, Acuña, Cifuentes-Muñoz et al. 2014).

**Fig. 4.**
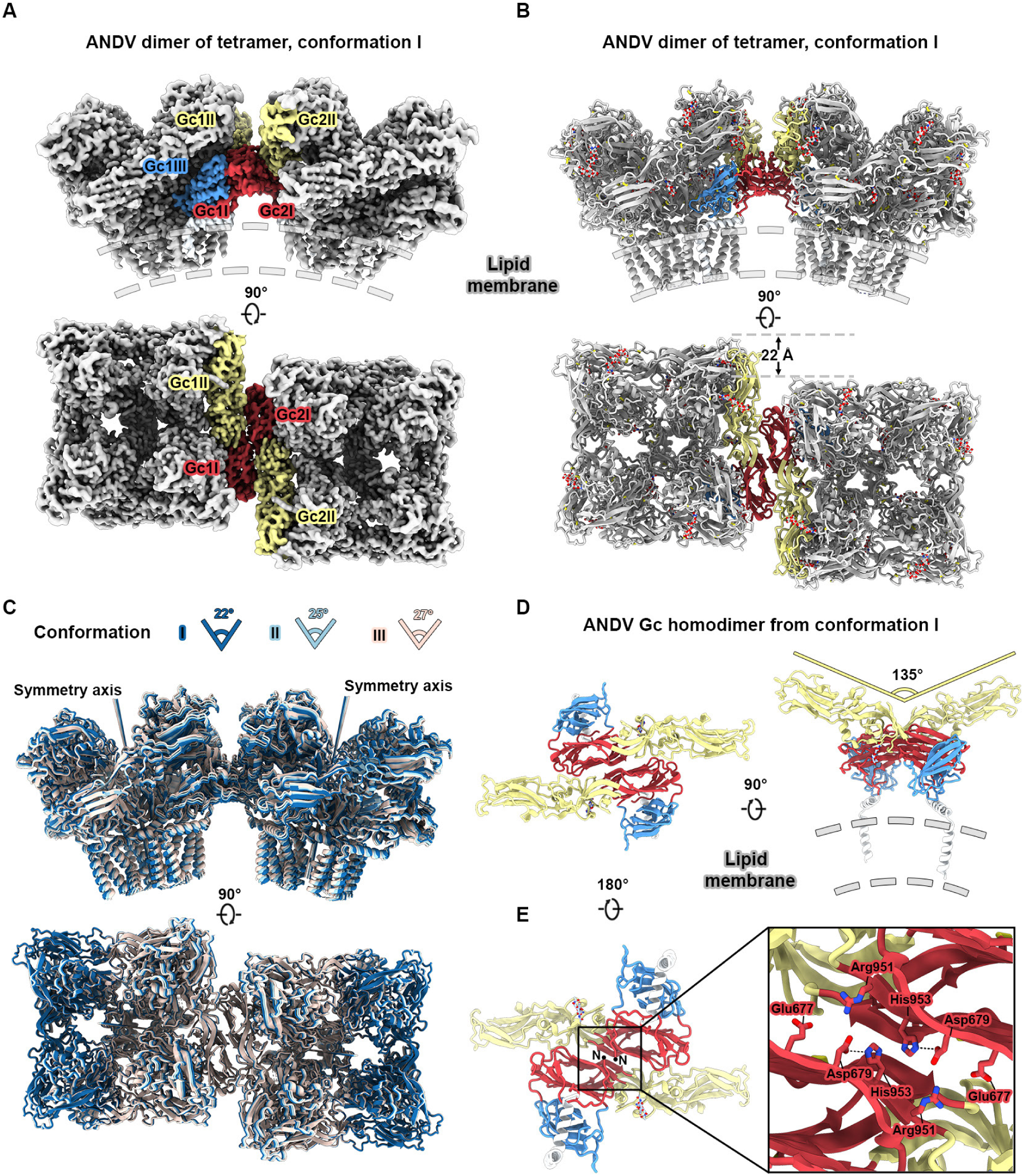
Organization and flexibility of dimers of ANDV Gn–Gc tetramers. (**A**) Cryo-EM map processed by EMReady of the ANDV dimer of tetramers, with regions of the interfacing Gc protomers colored by domain. (**B**) Structure of the ANDV dimer of tetramers shown as ribbons, colored as in (A). N-linked glycans are shown as sticks. (**C**) Alignment of the three resolved conformations of the ANDV dimer of tetramers, colored by conformation identity. The angle between the C4 symmetry axes for each conformation is noted. (**D**) Top and side views of the interfacing ANDV Gc dimers, shown as ribbons and colored by domain. (**E**) Bottom-up view of the interfacing ANDV Gc dimers. A zoomed-in view of the dimer interface shows interface sidechain as sticks. In all panels, nitrogen atoms are colored blue, oxygen atoms are red, and sulfur atoms are yellow. Hydrogen bonds are shown as black dashed lines.

**Table S1.**
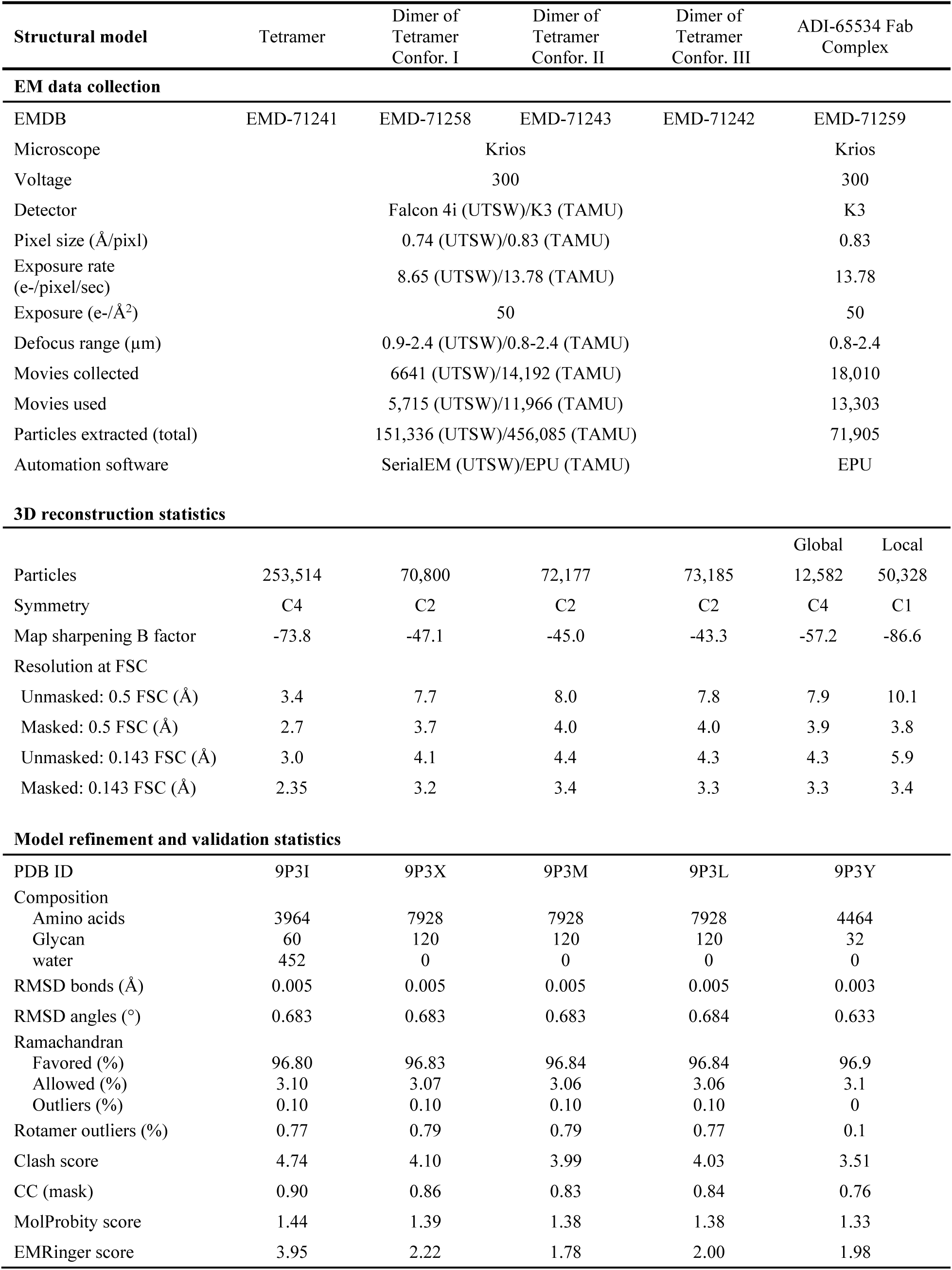
EM data collection and refinement statistics.

Inspection of the ANDV dimer-of-tetramers structures revealed that inter-tetramer association is mediated primarily by homotypic interactions between Gc protomers from adjacent tetramers (**Fig. 4B, S6,**), consistent with previously proposed models of hantavirus glycoprotein lattice organization (Hepojoki, Strandin et al. 2010, Bignon, Albornoz et al. 2019, Serris, Stass et al. 2020). Structural analysis of the Gc–Gc interfaces across the three conformational states revealed modest contact surfaces, with buried surface areas of 490 Å^2^, 454 Å^2^, and 494 Å^2^ for conformations I, II, and III, respectively, and an inter-protomer angle of approximately 135° (**Fig. 4D, S7**). These values are similar to the buried surface area of 547 Å^2^ observed in the crystallographic Gc dimer of HTNV (PDB ID: 5LJY), suggesting a conserved mode of Gc-mediated dimerization in hantaviruses (**Fig. S8**) (Bignon, Albornoz et al. 2019).

Primarily polar and electrostatic interactions were observed in the ANDV Gc homodimer interfaces, involving the sidechains of residues His953, Glu677, Asp679, and Arg951 (**Fig. 4E**). These are consistent with the interactions observed in the crystallographic HTNV Gc dimer, as expected given the strict conservation of these residues (Bignon, Albornoz et al. 2019). These interactions also agree well with prior mutagenesis studies, which demonstrated that mutation of Gc residues Glu677, Asp679, Arg951, and His953 in ANDV abolishes the fusion process in liposomes and transfected cells (Bignon, Albornoz et al. 2019). Notably, the limited inter-tetramer interface and centrally positioned His953 residues from opposing protomers create another pH-responsive assembly. Upon acidification, protonation of His953^Gc^ is expected to introduce electrostatic repulsion between the closely apposed imidazole rings, promoting dissociation of the Gc homodimer and disassembly of the glycoprotein lattice.

### ANDV VLPs enable determination of epitope- and lattice-level interactions of ADI-65534

Single-particle cryo-EM studies of antibodies in complex with ANDV-DM-eVLPs could be a useful approach for defining antibody–antigen interfaces and interactions with the glycoprotein lattice. To assess the feasibility of this approach, we first prepared cryo-EM grids of ANDV-DM-eVLPs complexed with ADI-65534 IgG—an affinity-matured, pan-hantavirus Gn– Gc-specific antibody derived from ADI-42898—which, as an IgG, was proposed to bind adjacent tetramers in the hantavirus lattice (Mittler, Serris et al. 2023). However, we were unable to produce grids containing separated VLPs, presumably due to cross-linking between VLPs by ADI-65534 IgG. We next attempted to complex the VLPs with ADI-65534 Fab and successfully obtained grids containing separated, Fab-bound VLPs. During particle curation, 2D class averages revealed adjacent Fab-bound tetramers (**Fig. 5A and S9**). Following refinement, a final stack of 12,582 particles picked from 13,303 accepted movies yielded a 3.3 Å global resolution reconstruction of a Fab-bound tetramer. Due to the reduced local resolution at the glycoprotein-Fab interface, local refinement was performed, resulting in a 3.3 Å resolution reconstruction of this region. **(Fig. 5B, S9, S10; Table S1)**.

**Fig. 5.**
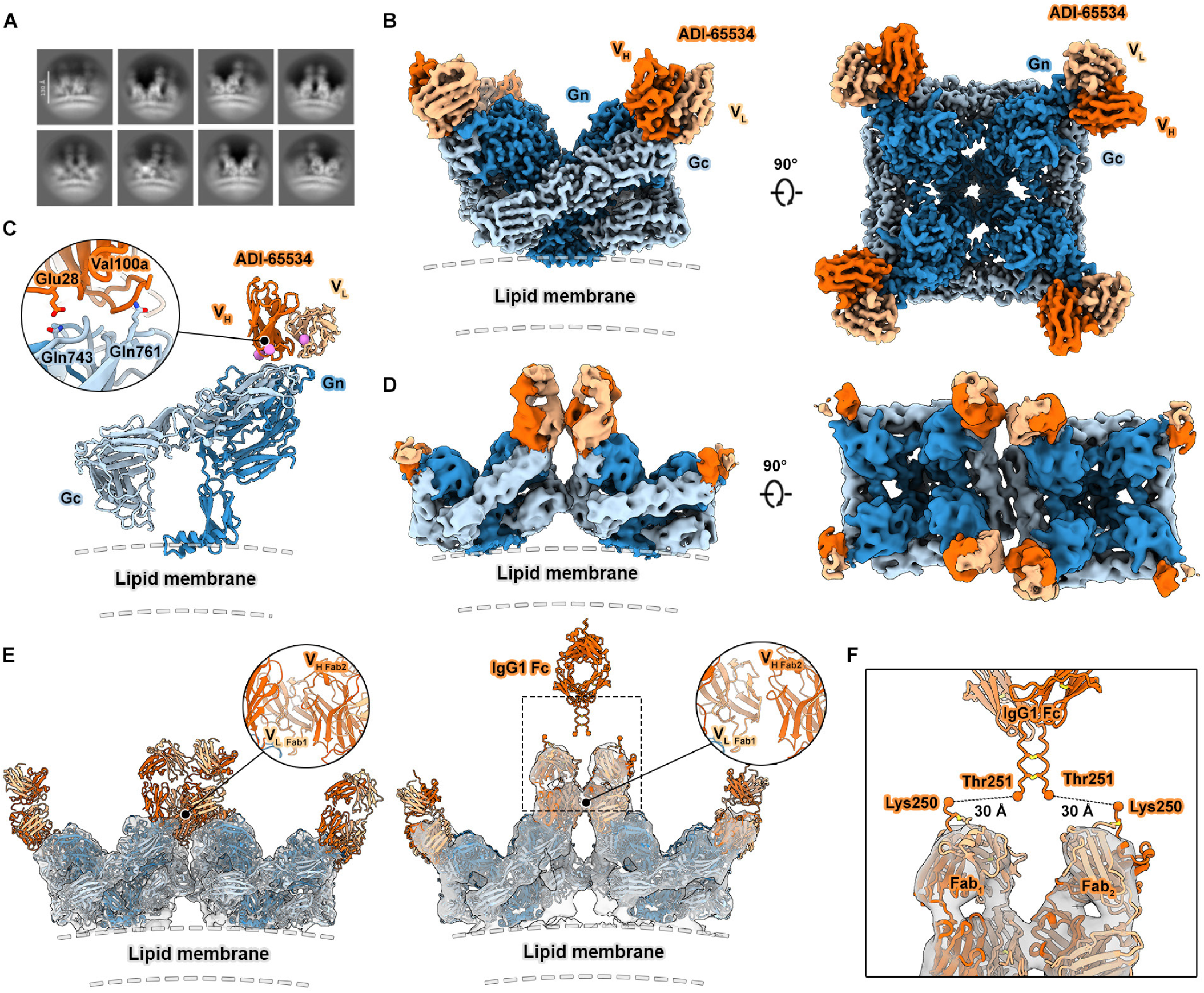
ANDV eVLPs enable determination of the epitope and lattice-level interactions of antibody ADI-65534. (**A**) Cryo-EM 2D class averages of ADI-65534 Fab-bound to ANDV glycoproteins on the surface of eVLPs. ANDV Gn– Gc tetramers. Scale bar: 130 Å. (**B**) A composite cryo-EM map processed by EMReady of the ANDV Gn-Gc tetramer with four ADI-65534 Fabs bound. (**C**) Interface between ADI-65534 Fab and the ANDV Gn–Gc heterodimer, shown as ribbons. The three substitutions in ADI-65534 relative to antibody ADI-42898 are shown as pink spheres. Select interface residues are shown as sticks. Nitrogen atoms are colored blue and oxygen atoms are red. (**D**) Side (*left*) and top-down (*right*) views of the dimer of tetramers map with eight ADI-65534 Fabs bound. (**E**) Modelling of the ADI-65534 Fab onto conformation I of the ANDV dimer of tetramers results in clashes between neighboring Fabs from adjacent tetramers. The in situ structure of the ANDV dimer of tetramers with eight Fabs bound is shown to the right. Enlarged views of the Fab-Fab interfaces are shown. An IgG1 Fc model predicted by AlphaFold3 is shown above two adjacent Fabs on neighboring tetramers. (**F**) Zoomed-in view of the hinge region of IgG1 Fc and ADI-65534 Fab, with the distance between the terminal residues of the Fab and Fc regions (orange spheres) indicated by dashed black lines.

ADI-65534 has only three substitutions relative to ADI-42898: T28E, G100aV (Kabat numbering), and C32V. Of these, only T28E and G100aV are in the paratope (**Fig. 5C, S10D**). It was previously determined that these substitutions result in increased ADI-65534 binding affinity to ANDV Gn–Gc and decreased affinity to PUUV Gn–Gc at acidic pH. The ADI-65534 Fab-bound ANDV glycoprotein tetramer model shows that the G100aV substitution leads to a favorable Van der Waals interaction with ANDV Gln761. The map quality for Glu28 was insufficient to confirm placement of the sidechain, however the Cα position of this residue supports several rotamer conformations that could enable a hydrogen bond with ANDV Gln743, which would not form with the shorter Thr28 found in ADI-42898 (Mittler, Serris et al. 2023).

Together, these data validate the biochemical basis for the increased affinity of ADI-65534 for ANDV Gn–Gc. Furthermore, we propose that substituting Gln for Glu28 may maintain hydrogen bonding with ANDV Gln743 and reduce electrostatic repulsion with PUUV Glu750, therefore improving binding to PUUV.

To assess whether a single ADI-65534 IgG molecule can simultaneously bind adjacent tetramers, a 6.8 Å resolution reconstruction of a dimer of Fab-bound tetramers was generated from 20,080 particles (**Fig. 5D, S9**). In this reconstruction, ADI-65534 Fab occupied all eight binding sites at the corners of the two tetramers, and two copies of the Fab-bound tetramer model described above could be placed in the reconstruction with minimal steric clashes. Notably, only conformation III of the apo ANDV dimer-of-tetramers allowed for the placement of the same Fab-bound tetramer models with minimal steric clashes. AlphaFold3 was used to model the ADI-65534 Fab and Fc domains based on an IgG1 backbone. The equidistant placement of the Fc between the C-termini of the heavy chains of adjacent Fabs showed an approximate 30 Å gap between residues Lys250 and Thr251, which connect the Fab and Fc regions (**Fig. 5E-F**). This 30 Å gap indicates that a single IgG1 molecule is constrained from bridging adjacent tetramers. However, IgG3, which has a hinge region approximately four times longer than that of the other subclasses, may possess sufficient flexibility to achieve this configuration. Thus, engineering ADI-65534 in an IgG3 backbone may enable bivalent binding of adjacent tetramers, thereby enhancing avidity and potentially increasing neutralization potency.

### LION^TM^/repRNA ANDV VLP vaccine candidates induce Gn–Gc-binding and rVSV-ANDV-neutralizing antibody titers in mice

To evaluate the immunogenicity of ANDV GPC variants, we synthesized sequences encoding the wildtype (WT) GPC (repGPC-WT), WT GPC with an eVLP tag (repGPC-WT-eVLP), the double modification (DM) GPC (repGPC-DM), and DM GPC with the eVLP tag (repGPC-DM-eVLP) into a plasmid DNA that encodes a self-amplifying replicon RNA (repRNA). In addition to the ANDV GPC open reading frame, repRNA encodes the Venezuelan equine encephalitis virus (VEEV), strain TC-83, replication machinery, which amplifies the antigen-encoding subgenomic message in a target cell. When delivered with the cationic nanocarrier Lipid InOrganic Nanoparticle (LION™), this repRNA/LION platform can transfect muscle cells at the injection site of the vaccinee (Kimura, Leal et al. 2023) and has been shown to drive VLP production from cells transfected with repRNA encoding certain viral glycoprotein antigens (Erasmus, Khandhar et al. 2018, Warner, Archer et al. 2024). After producing the repRNA ANDV GPC plasmids, we generated repRNA, formulated the repRNAs with LION^TM^, and proceeded to evaluate these vaccines in mouse immunogenicity studies.

Female C57BL/6 mice (n = 10/group) were immunized intramuscularly on days 0 and 35 with either 1 µg or 10 µg of each formulation, as well as an irrelevant antigen repRNA control (**Fig. 6A**). Blood was collected on days 0, 14, 35, and 49 to evaluate ANDV Gn–Gc-specific antibody binding titers via ELISA and neutralizing antibody titers via rVSV-ANDV-based assays. On day 14, LION/repGPC-DM-eVLP at a 10 µg dose induced a rapid binding antibody response, with mice displaying significantly higher titers relative to repGPC-WT (p = 0.045) and repGPC-DM (p = 0.034), respectively (**Fig. 6B**). The trend was similar, though not significant, at the 1 µg dose (**Fig. 6C**). By day 35, repGPC-DM-eVLP appeared to induce better binding antibody titers than all other variants tested at the 10 µg dose (repGPC-WT = 0.029, repGPC-DM = 0.001, repGPC-WT-eVLP = 0.006), with no significant differences at the 1 µg dose.

**Fig. 6.**
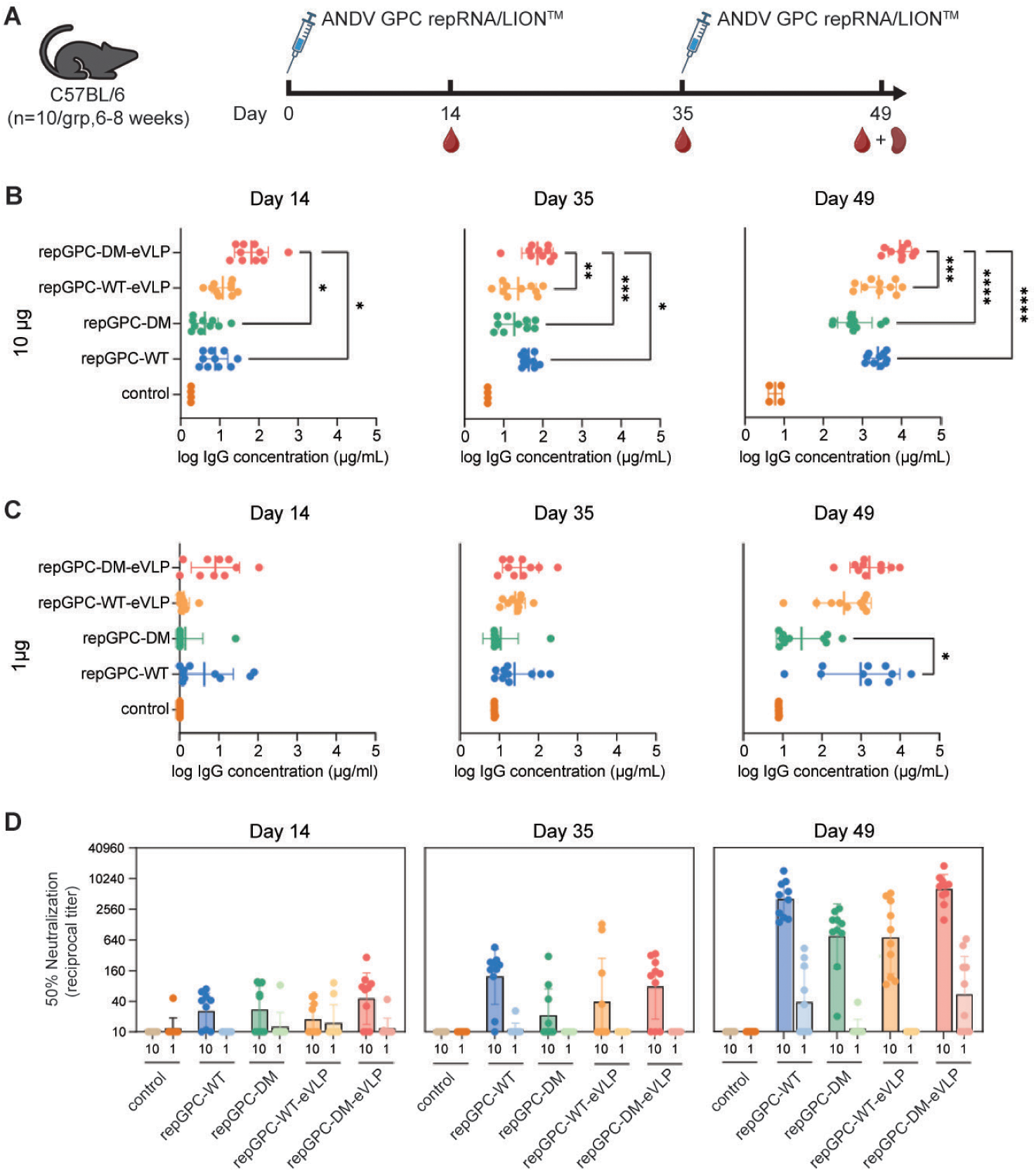
Immunogenicity of repRNA/LION^TM^ ANDV GPC constructs in C57BL/6 mice. (**A**) Schedule of immunizations and tissue collection for repRNA/LION^TM^ ANDV GPC vaccination studies. Female C57BL/6 mice (n = 10 per group, 6–8 weeks old) were immunized intramuscularly on Day 0 with either 1 µg or 10 µg of each construct. On Day 35, mice were boosted with the equivalent dose of the respective repRNA/LION^TM^ construct via intramuscular injection. Blood was collected on Days 14, 35, and 49. At study termination on Day 49, splenocytes were harvested for analysis. (**B**) Serum IgG concentrations of ANDV Gn– Gc-specific binding antibodies on Days 14 (left), 35 (middle), and 49 (right) following immunization of mice with the 10 µg dose. (**C**) Same as (B), but for the 1 µg dose. (**D**) Serum 50% neutralization reciprocal titers of rVSV-ANDV-neutralizing antibodies on Days 14 (left), 35 (middle), and 49 (right) following immunization of mice with 1 µg or 10 µg of each construct. For panels B-C, data are presented as the geometric mean with geometric standard deviation. Statistical significance was assessed using an ordinary one-way ANOVA with Tukey’s multiple comparisons test. *P < 0.05, **P < 0.01, ***P < 0.001, ****P < 0.0001.

Notably, mice receiving 10 µg of repGPC-DM-eVLP demonstrated significantly higher (p = 0.006) binding titers than mice receiving 10 µg of repGPC-WT-eVLP (**Fig. 6B**). At day 49, the repGPC-DM-eVLP construct continued to outperform all tested GPC variants, with higher binding antibody titers (repGPC-WT <0.0001, repGPC-DM = <0.0001, repGPC-WT-eVLP = 0.0009) at the 10 µg dose, but with no significant differences at the 1µg dose, except for inferior responses with repGPC-DM (p = 0.0416).

In terms of neutralizing antibody responses, we observed a dose dependence with low to no neutralizing antibody detected in all 1 µg dose groups, apart from repGPC-WT and repGPC-DM-eVLP groups, which received 2 doses of 1 µg, where 5 and 6 out of 10 animals, respectively, exhibited neutralizing antibodies at day 49 (**Fig. 6D**). At the 10 µg dose level, all vaccine candidates induced partial neutralizing responses by day 14 with only the repGPC-WT and repGPC-DM-eVLP groups achieving significantly higher geometric mean neutralizing titers (GMNTs) compared to the irrelevant control group 35 days after prime immunization (**Fig. 6D**). By day 49, two weeks after the boost immunization, the 10 µg repGPC-WT and repGPC-DM-eVLP vaccines elicited the highest GMNTs of 1:3,975 and 1:6,267, respectively, significantly higher than the 10 µg repGPC-DM and repGPC-WT-eVLP vaccines, which elicited GMNTs of 1:676 and 1:707, respectively (**Fig. 6D**). There were no statistically significant differences in T cell responses by ELISpot analysis, either in the 1 µg or 10 µg dose between repGPC-WT and repGPC-DM-eVLP (**Fig. S11**). Overall, these data suggest a plausible role for the eVLP tag, where the greatly increased in vitro expression observed from the eVLP-containing antigen compositions (**Fig. 1E**) can, to some extent, improve binding antibody responses in mice in a context-dependent manner but may not be sufficient to improve neutralizing antibody titers above that of the ANDV-WT GPC antigen composition in this vaccine modality.

## DISCUSSION

We initiated this study to determine high-resolution structures of the hantavirus glycoprotein tetramer and its homotypic interactions that underlie the pleomorphic lattice observed on virions. To avoid BSL-3 containment requirements associated with authentic ANDV, we produced and imaged rVSVs, native VLPs, and eVLPs, the latter created by the addition of an eVLP tag (EPM+EABR, (Hoffmann, Yang et al. 2023)) to the C-terminus of the ANDV M segment. We developed a single-particle cryo-EM data processing pipeline for lattice-forming glycoproteins on VLP membranes and determined high-resolution in situ structures of the ANDV Gn–Gc tetramer as well as a dimer of tetramers in three distinct but closely related conformations. This platform enabled structural analysis of antibody ADI-65534 in complex with ANDV-DM-eVLPs. Furthermore, we also explored the potential of ANDV eVLPs as a vaccine candidate expressed from repRNA via murine immunization.

The localization of the ANDV and CCHFV glycoproteins to the tips of rVSVs that we observed contrasts with the distribution of SARS-CoV-2 spikes on rVSVs and with native VSV G (Wu, Zhao et al. 2020, Zhou, Si et al. 2022, Xin, Kurien et al. 2025). It is possible that lateral forces generated within the lattice impose a curvature that is structurally compatible with the conical geometry at the virion tips rather than the cylindrical midsection of the VSV particle.

Such interactions are necessary to drive membrane curvature during budding of the virions because ANDV and CCHFV, unlike SARS-CoV-2 and VSV, do not encode matrix proteins (Plyusnin, Vapalahti et al. 1996, Deyde, Khristova et al. 2006). A second explanation for the polarized distribution is functional redundancy and steric hindrance between the Gn endodomain and the VSV matrix protein. In ANDV and CCHFV, the Gn endodomain acts as a surrogate matrix protein, tethering the glycoproteins to the ribonucleoprotein complex (Hepojoki, Strandin et al. 2010, Estrada and De Guzman 2011). In contrast, the VSV matrix protein forms a tightly packed scaffold around the nucleocapsid and interfaces with the inner leaflet of the viral membrane (Liu, Cao et al. 2021, Jenni, Horwitz et al. 2022, Zhou, Si et al. 2022). The compactness of the matrix layer along the cylindrical trunk may sterically exclude incorporation of Gn endodomains, whereas the comparatively looser matrix structure at the virion tips may accommodate or even be displaced by these domains. Further studies will be needed to determine whether bunyaviruses in the families *Peribunyaviridae* and *Phenuiviridae* exhibit a similar preferential localization, given that their glycoproteins also form a lattice (Martin, Lindsey-Regnery et al. 1985). Finally, the polarization of bunyavirus glycoproteins to the tips of rVSVs may have implications for the use of rVSVs as vaccines, as they may expose more lateral surfaces of the glycoproteins than found on the authentic viruses. Consistent with this idea, previous work indicates that rVSVs are more susceptible to neutralization by Gc-specific mAbs than their authentic hantavirus counterparts (Engdahl, Kuzmina et al. 2021, Mittler, Wec et al. 2022).

Structural analyses of pleomorphic viruses with glycoprotein lattices have traditionally relied on cryo-ET combined with sub-tomogram averaging (Li, Rissanen et al. 2016, Punch, Hover et al. 2018, Serris, Stass et al. 2020, Hover, Charlton et al. 2023). While these approaches have provided valuable insights into the architecture of glycoprotein lattices and the molecular mechanisms of entry, achieving near-atomic resolution has remained a challenge (Ni, Frosio et al. 2022, Liu, Zhou et al. 2023, Burt, Toader et al. 2024). Compounding these difficulties is the need for high-containment biosafety facilities when working with highly pathogenic viruses. In this study, we established a single-particle-based workflow capable of resolving glycoprotein lattices in a membrane environment to near-atomic resolution using eVLPs. This strategy circumvents biosafety limitations by enabling structure determination under BSL-2 containment conditions. Given the shared pleomorphic morphology among most bunyaviruses and the broad utility of the non-selective eVLP tag (Hoffmann, Yang et al. 2023), our platform provides a robust and adaptable framework for structural studies of lattice-forming viral glycoproteins and their immune complexes.

The high-resolution in situ ANDV glycoprotein tetramer structures presented here reveal multiple stabilizing interfaces that preserve the tetramer in its prefusion conformation while allowing for pH-dependent activation. Several of these, including the Gn^B^2–Gc1III interface and the Gn^H^2–Gc1 interface, are described here for the first time. This elaborate network of interactions likely accounts for the ability of the ANDV glycoprotein lattice on rVSVs and VLPs to survive purification, whereas the SARS-CoV-2 S protein was found to exhibit a predominantly post-fusion conformation on the surface of rVSV virions. The conservation of residues at these interfaces across hantaviruses (**Fig. S8**) supports a shared mechanism underpinning environmental resilience, consistent with the virus’s ability to remain infectious for days outside the host (Kallio, Klingström et al. 2006, Hardestam, Simon et al. 2007) and to transmit via aerosolized excreta (Jonsson, Figueiredo et al. 2010, Avšič-Županc, Saksida et al. 2019, Vial, Ferrés et al. 2023). The structures presented here and previously (Serris, Stass et al. 2020) suggest that to enable fusion, ANDV exploits pH-sensitive triggers by embedding one or more histidine residues at the center of these stabilizing interfaces. Upon acidification, protonation of these histidines introduces repulsive forces, either between neighboring positive charges or adjacent protonated histidines, destabilizing the interfaces and initiating conformational rearrangements, akin to the mechanism observed in influenza virus hemagglutinin (Stevens, Corper et al. 2004, Mair, Meyer et al. 2014), flavivirus envelope protein E (Bressanelli, Stiasny et al. 2004, Fritz, Stiasny et al. 2008), and VSV G protein (Rücker, Wieninger et al. 2012, Beilstein, Abou Hamdan et al. 2020).

The LION^TM^/repRNA immunogenicity studies demonstrated that, both for neutralizing and Gn–Gc-specific antibody responses, the inclusion of an eVLP tag in the WT GPC sequence resulted in a reduced immune response. However, the inclusion of the two substitutions (DM) with an eVLP tag may rescue the immune response, likely by driving better expression to the plasma membrane. That ANDV-DM-eVLP consistently drove higher binding antibody titers— but not neutralizing antibody titers—relative to ANDV-WT across all time points merits further study. This finding stands in contrast to studies of SARS-CoV-2 spike-eVLPs (S-eVLP), where mRNA encoding S-eVLPs consistently induced higher neutralizing antibody titers across all time points compared to spike mRNA alone when 2 μg S-eVLP mRNA encapsulated in lipid nanoparticles was administered by similar intramuscular injections (Hoffmann, Yang et al. 2023). Thus, the relationship between eVLP-glycoprotein fusions and anti-viral neutralizing antibody responses in mice may be glycoprotein-or eVLP-repRNA-dependent. Further immunogenicity studies will be needed to address this question and identify the optimal hantavirus vaccine antigen, including structure-based design efforts involving stabilization of tetramer interfaces and display of stabilized tetramer ectodomains onto soluble scaffolds, which will now be enabled by the high-resolution structures presented here.

## METHODS

### Recombinant vesicular stomatitis virus production, purification and fixation

A recombinant VSV expressing CCHFV GPC was generated by plasmid-based rescue as described (Dieterle, Haslwanter et al. 2020), plaque-isolated, and sequenced. rVSV-CCHFV, rVSV-ANDV, (Kleinfelter, Jangra et al. 2015) and rVSV-SARS-CoV-2 (Dieterle, Haslwanter et al. 2020) were propagated on HUH7.5 cells for large-scale virus production. Two days post-infection, the supernatants were isolated and clarified by centrifugation at 4000 × g for 10 min. Virions in the cleared supernatants were pelleted by ultracentrifugation in an SW28 rotor (Beckman-Coulter Life Sciences) at 20,000 rpm. Subsequently, viral pellets were resuspended in PBS and fixed with 0.2 mM glutaraldehyde for 1 h on ice. The fixative was then quenched with 50 mM Tris-Cl (pH 7.5). The viral preparations were stored at -80 °C prior to downstream structural analysis.

### ANDV VLP expression and purification

The codon-optimized nucleic acid sequence of the ANDV M segment (Strain: Chile-9717869. GenBank: NC_003467.2) was synthesized by GenScript and cloned into the pCAG vector under the control of the chicken β-actin promoter, incorporating an in-frame 8×His tag to generate the ANDV-WT construct. To generate the eVLP-tagged construct (ANDV-WT-eVLP), codon-optimized sequences encoding the eVLP tag (EPM+EABR) (Hoffmann, Yang et al. 2023) and an 8×His tag were synthesized and inserted in place of the 8×His tag in the ANDV-WT construct.

Site-directed mutagenesis was then used to introduce the double mutations V535K and S1096L into both the ANDV-WT and ANDV-WT-eVLP constructs, yielding ANDV-DM and ANDV-DM-eVLP, respectively.

Plasmids encoding ANDV constructs were transfected into FreeStyle^™^ 293-F cells (Thermo Fisher Scientific) at a density of 1 × 10^⁶^ cells/mL using 25 kDa polyethyleneimine (PEI; PolySciences). Transfected cells were cultured in suspension at 37 °C in a shaking incubator with 8% CO_₂_ and 80% humidity for 72 hours. Supernatants were harvested by centrifugation, passed through 0.22 μm Rapid-Flow^™^ filters (Nalgene), and concentrated to ∼10 mL using a Centricon® Plus-70 filter unit (MilliporeSigma). The concentrated supernatant was layered onto a discontinuous sucrose gradient (20%, 50%, and 70% sucrose in PBS) and subjected to ultracentrifugation at 135,000 × g for 2 hours at 4 °C. Fractions corresponding to VLPs were collected, pooled, and further concentrated to ∼60 μL in PBS (pH 7.4) for cryo-EM grid preparation.

To evaluate expression levels among the ANDV-VLP constructs, supernatants from 200 mL cell cultures transfected with different constructs were collected, filtered, and concentrated to the same volume of 4 mL. Then, the aliquoted and concentrated supernatants were analyzed by Western blot using monoclonal antibodies against the His tag (Thermo Fisher Scientific, Cat. # MA1-21315) followed by IRDye® 800CW goat anti-mouse IgG secondary antibody (LI-COR Biosciences, Cat. # 926-32210). Protein bands on PVDF membranes were visualized using the LI-COR Odyssey imaging system, and band intensities were quantified using the accompanying software after background subtraction.

### ADI-65534 Fab expression and purification

Codon-optimized nucleotide sequences encoding the variable regions of ADI-65534 (Mittler, Serris et al. 2023) were synthesized and cloned by GenScript into expression plasmids encoding human IgG1 heavy and light chain constant domains. A human rhinovirus 3C (HRV3C) protease cleavage site was incorporated into the linker region to facilitate digestion into Fabs. Heavy and light chain plasmids were co-transfected into FreeStyle^™^ 293-F cells (Thermo Fisher Scientific) at a density of 1 × 10^⁶^ cells/mL using 25 kDa polyethyleneimine (PEI; PolySciences). Transfected cultures were incubated for 5 days at 37 °C in a shaking incubator maintained at 8% CO_₂_ and 80% humidity. Following incubation, supernatants were harvested by centrifugation, filtered through 0.22 μm Rapid-Flow^™^ filters (Nalgene), and flowed over Pierce^™^ Protein A/G Plus agarose resin (Thermo Fisher Scientific). After washing the column with PBS, the IgG-bound resin was incubated with HRV3C protease (1:100 wt/wt) for 1 hour at room temperature. The flow-through containing ADI-65534 Fab was collected and concentrated using a 30 kDa MWCO filter, flash-frozen, and stored at -80 C.

### Cryo-EM grid preparation and data collection

Cryo-EM grids for ANDV-VLP samples were prepared using R1.2/1.3 300 mesh UltrAuFoil grids (Quantifoil). Grids were glow-discharged and plunge-frozen into liquid ethane using a Vitrobot Mark IV (Thermo Fisher Scientific) with a blot force of 4 and a blot time of 20 s. For the ADI-65534 Fab–ANDV-DM-eVLP complex, ADI-65534 Fab (1.4 mg/mL) was mixed with ANDV-DM-eVLP at a 3:1 volume ratio and incubated on ice for 3 min. The mixture was then applied to glow-discharged UltrAuFoil grids and vitrified under identical conditions. Initial screening of grids was performed on a 200 kV Glacios (Thermo Fisher Scientific) at The University of Texas at Austin. Grids exhibiting optimal ice thickness were transferred to the cryo-EM facilities at UT Southwestern Medical Center (UTSW) and Texas A&M University (TAMU) for high-resolution data collection.

The ANDV-DM-eVLP dataset collected at UTSW comprised 6,641 movies collected on a 300 kV Titan^™^ Krios (Thermo Fisher Scientific) equipped with a cold (cathode) field-emission gun (cold-FEG), Falcon 4i detector, and Selectris energy filter. Data were acquired at a physical pixel size of 0.74 Å, with a total dose of 50 e⁻/Å^2^ and a 10 eV energy filter slit width.

The second ANDV-DM-eVLP dataset, along with the 65534 Fab–ANDV-DM-eVLP dataset, were collected at TAMU on a Titan™ Krios G4 (Thermo Fisher Scientific) equipped with an X-FEG, K3 Summit camera (Gatan), and BioContinuum Imaging Filter (Gatan). A total of 14,192 movies were acquired at a pixel size of 0.832 Å, with a total dose of 50 e⁻/Å^2^ and a 15 eV energy filter slit width. Full data collection parameters are summarized in **Table S1**.

### EM data processing and model building

All datasets were initially imported into CryoSPARC v4.6.0 (Punjani, Rubinstein et al. 2017) for patch-based motion correction, contrast transfer function (CTF) estimation, and exposure curation. crYOLO v1.9.9 (Wagner, Merino et al. 2019) was used for particle picking, guided by a trained model based on manually picked particles from 100 motion-corrected, dose-weighted micrographs. The resulting particle coordinates were imported into CryoSPARC for extraction.

Particles were first extracted using a large box size and subjected to two rounds of 2D classification, *ab initio* reconstruction, heterogeneous refinement, and homogeneous refinement to generate a low-resolution map containing multiple tetramers. To isolate individual tetramers or dimer-of-tetramer particles, coordinates were recentered on the centroid of each subunit, followed by re-extraction using a smaller box size. These recentered particles were processed through additional rounds of 2D classification, *ab initio* reconstruction, heterogeneous refinement, homogeneous refinement, 3D classification, and reference-based motion correction to obtain the final high-resolution EM maps.

For combined processing of the UTSW and TAMU datasets of ANDV-DM-eVLP, each dataset was initially refined independently to high resolution. Relative pixel scaling factors were determined using ChimeraX (Meng, Goddard et al. 2023) following established protocols (Wilkinson, Kumar et al. 2019). Raw UTSW movies were then rescaled to match the TAMU dataset using the relion_image_handler command from RELION v5.0 (Burt, Toader et al. 2024), and the datasets were merged for combined processing in CryoSPARC. The full data processing workflow for ANDV-DM-eVLP is shown in **Figure S2** and **Figure S5**. The ADI-65534 Fab dataset was processed similarly (**Figure S9**), except initial models of the tetramer and dimer of tetramers were derived from the ANDV-DM-eVLP reconstructions.

An initial atomic model of the ANDV tetramer was built using ModelAngelo (Jamali, Käll et al. 2024) with amino acid sequences provided as input. Manual inspection, model adjustment, and glycan building were performed in Coot v0.9.8.92 (Emsley, Lohkamp et al. 2010), followed by iterative rounds of real-space refinement in Phenix v1.21.2 (Liebschner, Afonine et al. 2019). Glycan structure validation was performed using the Privateer web server (Dialpuri, Bagdonas et al. 2024). Water molecules were placed using Douse within Phenix and manually curated in Coot. A final round of refinement in Phenix yielded the complete atomic model.

For model building of the ANDV dimer of tetramers, the refined tetramer model was docked into the dimer maps and refined using one round of real-space refinement in Phenix. To construct the ANDV-65534 Fab complex model, the refined tetramer model (excluding water molecules) was combined with the scFv of ADI-42898 from PDB ID:7QQB. Mutations were introduced in Coot to convert the Fab to the ADI-65534 sequence, and the complex was subsequently refined in Phenix. The final model represents the ANDV tetramer bound by four Fab molecules. Refinement statistics for all models are summarized in **Table S1**.

### Structural model analysis

All structural analyses were performed in ChimeraX (Meng, Goddard et al. 2023) for the ANDV-DM-VLP tetramer and dimer-of-tetramer model. To visualize unmodeled densities, a difference map was computed using TEMPy-DiffMap (Joseph, Lagerstedt et al. 2020) within the CCP-EM suite (Burnley, Palmer et al. 2017). Briefly, the experimental EM map was thresholded at a value of 0.01 and filtered using a dust filter to remove noise. A simulated map was then generated from the atomic model, and the difference between the filtered experimental map and the simulated map was calculated. To isolate regions of interest, the resulting difference map was segmented using the Segger tool (Pintilie, Zhang et al. 2010)in ChimeraX, and irrelevant densities were manually subtracted.

To model the dimer of Fab-bound tetramers, two ADI-65534-ANDV-DM-eVLP models were fit into the Fab-bound dimer-of-tetramer map. To model the constant regions of the antibody, two separate AlphaFold3 (Abramson, Adler et al. 2024) models were generated for the full ADI-65534 Fab and Fc regions. The Fab was aligned to the model by the variable fragments using the MatchMaker command in ChimeraX. The Fc was manually placed between two Fabs from adjacent tetramers, and the distance between the C-terminus of the Fab heavy chain and the N-terminus of the Fc was measured using ChimeraX.

### Sequence conservation analysis

The ANDV M segment amino acid sequence (NP_604472.1) was used as a seed to search against Uniprot database using Hmmer webserver (Potter, Luciani et al. 2018) for similar sequences. The hits were manually thresholded based on the species, hit positions and bit scores. To avoid potential bias against certain species, 10 sequences with the highest bit scores from ANDV, Sin Nombre virus (SNV), Hantaan virus (HTNV), Seoul virus (SEOV), Puumala virus (PUUV), Dobrava-Belgrade virus (DOBV) and Tula virus (TULV) were selected and aligned using Clustal Omega (Madeira, Madhusoodanan et al. 2024). The alignment was then used to color the ANDV tetramer surface in ChimeraX, highlighting the conservation of each residue.

### Animals, biosafety and ethics

All animal work was performed in strict accordance with the recommendations described in the Guide for the Care and Use of Laboratory Animals of the Office of Animal Welfare, National Institutes of Health and the Animal Welfare Act of the US Department of Agriculture, in an AAALAC-accredited facility (Assurance ID: D16-00496) and study protocol approved by the Charles River Accelerated Development Laboratory’s (CRADL) Institutional Animal Care and Use Committee IACUC (Protocol #: EB24-400-400). Animals were housed under controlled conditions of humidity, temperature and light (12-h light/12-h dark cycles). Food and water were provided *ad libitum*.

### Synthesis and characterization of RNA for ANDV GPC variants

Double-stranded DNA fragments encoding ANDV GPC variants were synthesized and cloned into a plasmid backbone between the PflFI and SacII restriction enzyme sites. This backbone harbors the 5′ and 3′ untranslated regions and nonstructural open reading frame of Venezuelan equine encephalitis virus, strain TC-83, as well as a T7 DNA-dependent RNA polymerase promoter and kanamycin-resistance gene cassette. Plasmids were sequence-verified and subsequently linearized by digestion with the NotI restriction enzyme. RNA was then transcribed and capped from purified linear DNA templates by T7 RNA polymerase and Vaccinia virus capping enzyme reactions, as previously described (Warner, Archer et al. 2024). The resulting RNA was analyzed by capillary gel electrophoresis to confirm production of full-length transcripts.

### Mouse immunogenicity studies

Female C57BL/6 mice aged 6–8 weeks (*n* = 10/group) were sourced from Jackson Laboratories and acclimated prior to study initiation. Mice were immunized intramuscularly with 1 µg or 10 µg of repRNA/LION™ complexed at an N:P ratio of 15. The vaccine was delivered via a single 50 µL injection on a prime/boost schedule, with immunizations on Day 0 (prime) and Day 35 (boost). Blood samples were collected via submental vein bleed on days 0, 14, and 35, and via cardiac puncture on day 49. On the day of harvest, tissues were collected and stored in RPMI 1640 medium (Invitrogen) supplemented with 10% heat-inactivated fetal calf serum (Gibco) and stored on ice until processing into single-cell suspensions. Sham-immunized animals received an equivalent dose of repRNA encoding Rift Valley fever virus glycoprotein complexed with LION™ as an irrelevant antigen control.

### rVSV-ANDV production and neutralization assays

Replication-competent, rVSVs expressing an eGFP reporter and bearing ANDV Gn/Gc (NP_604472.1) were generated using a plasmid-based rescue system in 293T cells and propagated on Vero cells as described previously (Whelan, Ball et al. 1995, Kleinfelter, Jangra et al. 2015). The sequence of Gn/Gc was verified by amplifying viral genomic RNA by RT-PCR and analyzed using Sanger sequencing. For neutralization experiments, pre-titrated rVSV-ANDV particles, with a multiplicity of infection (MOI) of 0.1, were incubated with decreasing concentrations of heat-inactivated mouse sera (starting at 1:100 dilution) at room temperature (RT) for 1 h before being added to Vero cell monolayers in 96-well plates. 14–16 hours post-infection, cells were fixed with 4% formaldehyde (Sigma) and counter-stained with Hoechst nuclear stain (Invitrogen). Number of infected cells (GFP^+^ cell counts) and viral infectivity (%GFP^+^ cells) were measured by automated enumeration of eGFP-expressing cells from captured images using Cytation5 cell imaging multi-mode reader with Agilent Biotek Gen5

Microplate Reader and Imager software (BioTek, V.3.2). The infectivity was normalized to that obtained without sera and subjected to nonlinear regression analysis to derive neutralization titers (NT50) using a 4-parameter, variable slope sigmoidal dose-response (GraphPad Prism).

### ELISAs

High-Bind plates (Corning, Cat. #3700) were coated with recombinant ANDV-Gn/Gc protein produced by GenScript. The protein was expressed in TurboCHO cells and purified with a HisTrap^TM^ FF Crude column and a HiLoad 26/600 Superdex 200pg column. Protein purity was verified by SDS-PAGE analysis. ANDV Gn/Gc was diluted to a final concentration of 2 µg/mL and allowed to bind plates by incubating in phosphate-buffered saline (PBS) overnight at 4 °C. On the day of the assay, plates were blocked using 1% non-fat dry milk in 1× PBS containing 0.05% Tween-20 (PBS-T). Serial dilutions of mouse sera were prepared in blocking buffer, added to the plates, and incubated at room temperature for 1–2 h. Bound antibody was detected using anti-mouse IgG conjugated to horseradish peroxidase (Goat Anti-Mouse IgG(H+L)-HRP, Southern Biotech, Cat. #1036-05) at a final dilution of 1:10,000. Plates were developed with SureBlue Reserve^™^ 3,3′,5,5′-Tetramethylbenzidine (TMB) 1-Component Microwell Peroxidase Substrate (SeraCare, Cat. #5120-0077). The reaction was quenched with TMB Stop solution (SeraCare, Cat. #: 5150-0021), and absorbance was read at 450 nm. Endpoint titers were defined as the reciprocal serum dilution that yielded an absorbance value ≥2 standard deviations above background, determined by curve fitting and interpolation.

### IFN-γ ELISpot assay

Splenocytes were isolated from mice on day 49 post-prime immunization and passed through a 70 µm cell strainer (Corning, Cat. #431751). Multiscreen® 96-well plates containing a hydrophobic PVDF membrane (Millipore Sigma, Cat. #MIAPS4510) were coated with rat anti-mouse IFN-γ capture antibody (BD, Catalog #551216, Lot 4099272) at a final concentration of 10 µg/mL in 1X PBS overnight at 4 °C. The plates were washed using 1X PBS and then blocked for 2 hours at room temperature with RPMI 1640 medium (Invitrogen, Cat. #11875093) containing 10% heat-inactivated fetal calf serum (Invitrogen, Cat. #A5670701). Splenocytes were plated at 5 × 10^5^ cells/well and stimulated with peptide pools from a custom ANDV-GPC peptide library. Peptides (15-mers overlapping by 10 amino acids) were synthesized by GenScript, reconstituted in 90% DMSO/10% water at 1 mg/mL, and pooled in an approximately equimolar ratio with 15–20 peptides per pool. Pools were further combined into three mega-pools, each comprising peptides from five individual pools. All stimulations were performed at a final concentration of 1.5 µg/ml/peptide and cultured for 20 h (37 °C, 5% CO_2_). Negative control wells were incubated with an equivalent volume of DMSO, and positive control wells were stimulated with 20 ng/mL phorbol myristate acetate (PMA) and 1 μg/mL ionomycin. After incubation, biotinylated anti-mouse IFN-γ antibody (BD, Catalog #554410, Lot# 3166827 or 1313950) at a final concentration of 1 µg/mL and 1-Step™ NBT/BCIP Substrate Solution (Invitrogen, Catalog #34042, Lot #ZG395931) were used to detect IFN-γ-secreting cells. Spot-forming cells were enumerated using an Immunospot Analyzer from CTL Immunospot software (Cellular Technology Ltd).

### Data and code availability

Atomic coordinates and cryo-EM maps have been deposited in the PDB and EMDB, respectively, with accession codes: ANDV tetramer (PDB ID: 9P3I, EMD-71241), ANDV dimer of tetramers conformation I (PDB ID: 9P3X, EMD-71258), ANDV dimer of tetramers conformation II (PDB ID: 9P3M, EMD-71243), ANDV dimer of tetramers conformation III (PDB ID: 9P3L, EMD-71242), ANDV tetramer in complex with ADI-65534 Fab (PDB ID: 9P3Y, EMD-71259), and ANDV dimer of tetramers bound with ADI-65534 Fab (EMD-71260). This paper does not report original code.

## ACKNOWLEDGMENTS

We thank Mahtab Beikzadeh and James Guerra for their help with protein expression and other members of the McLellan Laboratory for providing helpful comments on the manuscript, particularly Nicole Johnson and Yuna Shin, as well as Justine Meccio for administrative support. We thank Allie Lafferty, Hathaipat Arayangkul, Brian Granger, Karen Gaffney, Chanel Estrada, Carrie Novak, Stephanie Park, Shibbu Sharma, Savannah Moritzky, Kianna Sinfuego and Samantha Randall for their technical assistance. We thank Axel Brilot and Evan Schwartz at the Sauer Structural Biology Laboratory at UT Austin for assistance with EM grid screening as well as Zhe Chen, Wei Liu, and Yang Li at the Structural Biology Lab at UT Southwestern Medical center (partially supported by CPRIT grant RP220582) and Gaya P. Yadav at the Laboratory for Biomolecular Structure and Dynamics at Texas A&M University for their help with data collection. This work was funded in part by NIH grants U19AI181977 (K.C., J.S.M., J.H.E.) and U19AI142777 (K.C., J.S.M.) as well as Welch Foundation grant number F-0003-19620604 (J.S.M.), Welch Foundation grant number F-2244-20250403 (Z.K.), and CPRIT grant RR230050 (Z.K.). We acknowledge The University of Texas College of Natural Sciences and CPRIT award RR160023 for support of the EM facility.

## AUTHOR CONTRIBUTIONS

Conceived the project: L.Q., J.H.E., K.C., J.S.M.

Designed and performed experiments: L.Q., E.Mc., M.M.S., E.T.S., J.B., E.Mi., K.H., T.H., H.S.K., Z.K., J.S.M.

Supervision: N.L.W., J.H.E, K.C., J.S.M.

Manuscript writing, first draft: L.Q., E.M., E.T.S., J.H.E., J.S.M. Manuscript review and editing: All authors

## DECLARATION OF INTERESTS

E. Mittler and K. Chandran are coinventors on a patent application describing antibodies against hantaviruses, including ADI-65534.

