## Supplemental Figures for "High-resolution in situ structures of hantavirus glycoprotein tetramers"

#### Figure S1

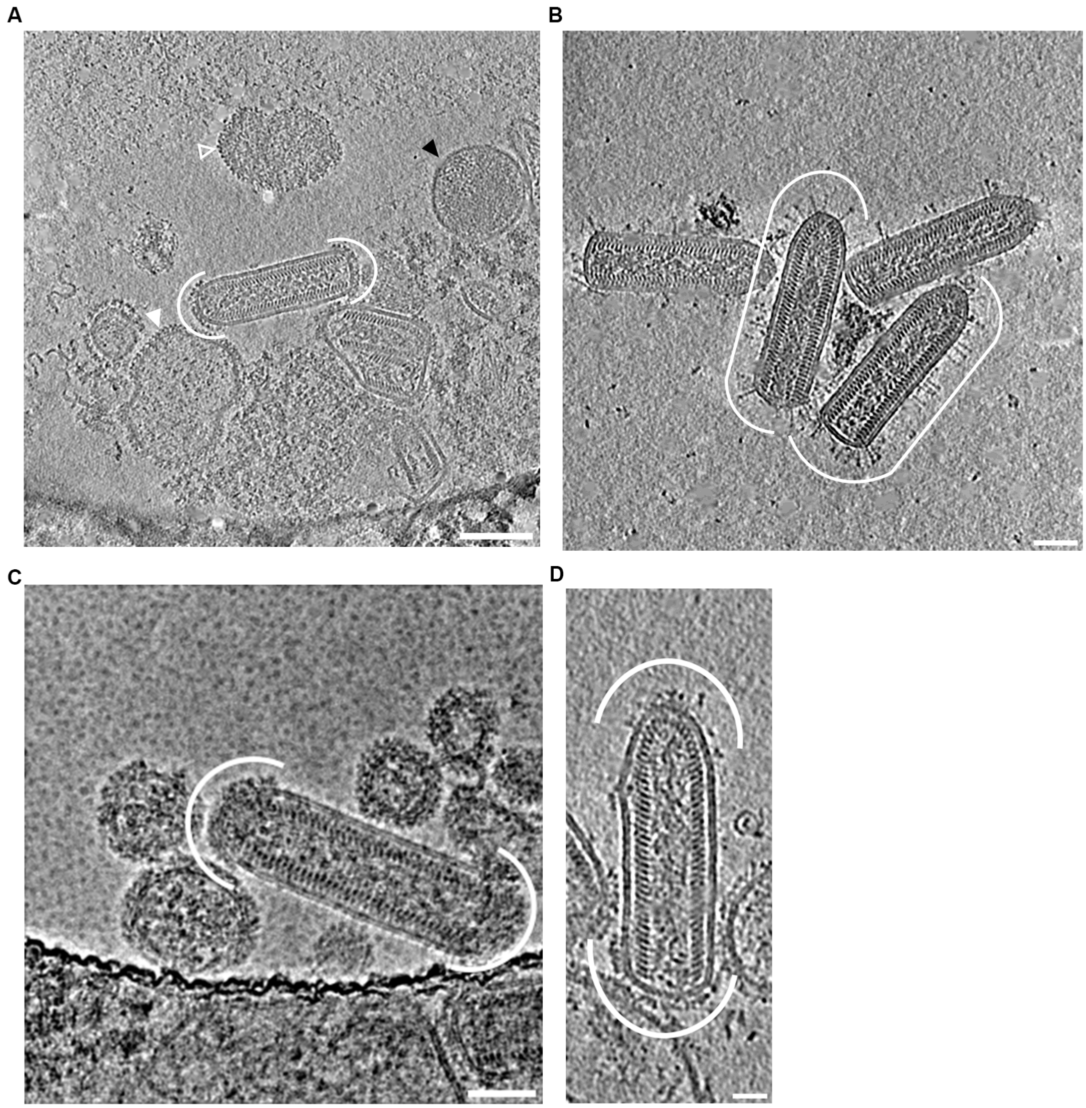


**Fig. S1. Tomography slices of rVSV-ANDV, rVSV-SARS-CoV-2 and rVSV-CCHFV**

(**A**) A 16 nm tomographic slice of rVSV-ANDV reveals a heterogeneous sample composition, including an intact rVSV-ANDV virion (white curved lines), an rVSV-ANDV virion with two broken nucleoprotein arrangements, a virus-like particle coated with ANDV glycoproteins (filled white arrowhead), a bald vesicle (black arrowhead), and a VLP displaying ANDV glycoproteins and unknown densities (white hollow arrowhead). (**B**) A 14 nm tomographic slice of rVSV-SARS-CoV-2 reveals several VSV virions with surfaces decorated with spike proteins. Scale bar, 50 nm. (**C**) Cryo-EM 2D projection image of an rVSV-CCHFV virion shows that the CCHFV glycoproteins also preferentially localize to both tips of the rVSV virion. Multiple VLPs displaying the CCHFV spike protein can also be observed. Scale bar, 50 nm. (**D**) A 16 nm tomographic slice of rVSV-CCHFV virion confirms the preference of CCHFV glycoproteins covering both tips of the virion. Scale bar, 25 nm.

#### Figure S2

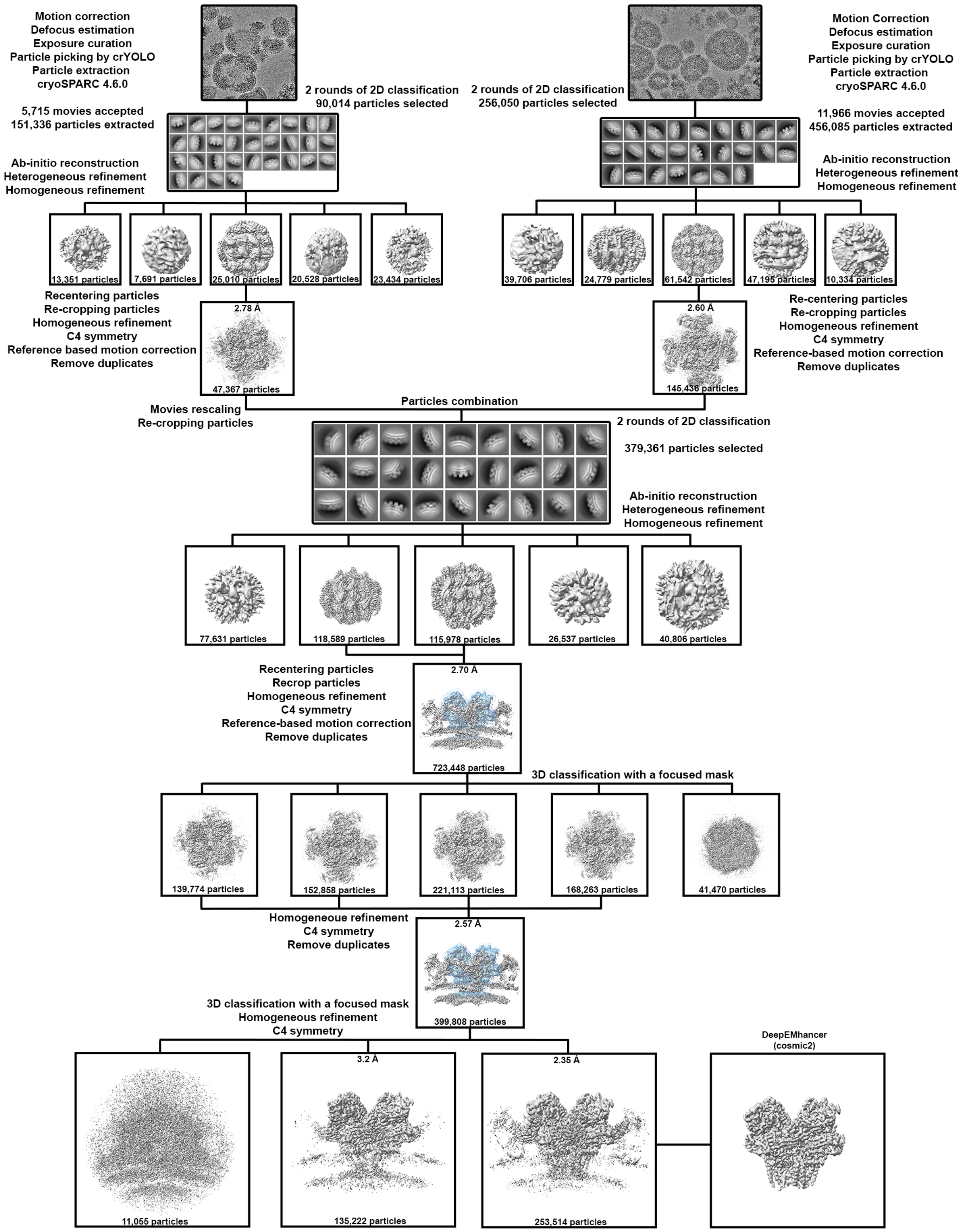


**Fig. S2. Cryo-EM data processing workflow for the ANDV Gn–Gc tetramer.**
Flowchart illustrating the cryo-EM data processing steps used to obtain the high-resolution structure of the ANDV Gn–Gc tetramer. Particle counts at each stage are indicated, along with the corresponding resolutions of the 3D reconstructions. Masks used for recentering and focused 3D classification are represented as blue transparent surfaces.

#### Figure S3

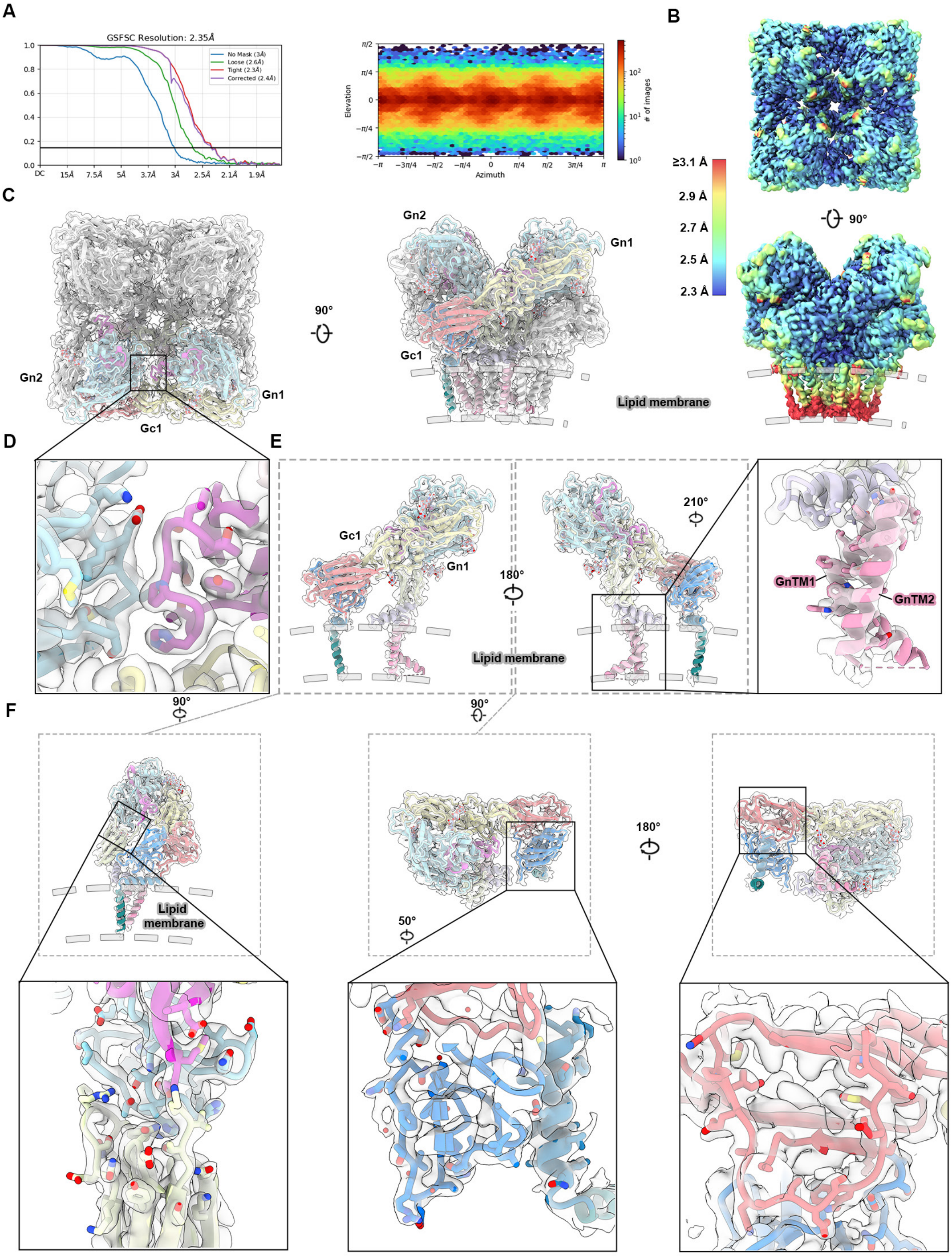


**Figure S3. Cryo-EM map validation and model fitting for the ANDV Gn–Gc tetramer.**
(**A**) Global gold-standard Fourier shell correlation (GSFSC) curve and particle orientation distribution. (**B**) Cryo-EM map of the ANDV Gn–Gc tetramer colored by local resolution. Densities from adjacent tetramers were masked out. The transmembrane domains are shown overlapping a lipid bilayer, indicated by gray dashed curves. (**C**) Overview of the map-to-model fit for the ANDV Gn–Gc tetramer. The model is shown as gray ribbons, with one Gn1–Gc1–Gn2 heterotrimer colored by domain (color scheme as in Fig. 3). (**D**) Close-up view of the $Gn^{H}1$–$Gn^{H}2$ interface and its fit to the EM map. (**E**) Map-to-model fit of the Gn1–Gc1 heterodimer, highlighting the kink in the GnTM1 helix. (**F**) The map-to-model fit exhibits clear densities for sidechains of residues that were absent in previous structures.

#### Figure S4

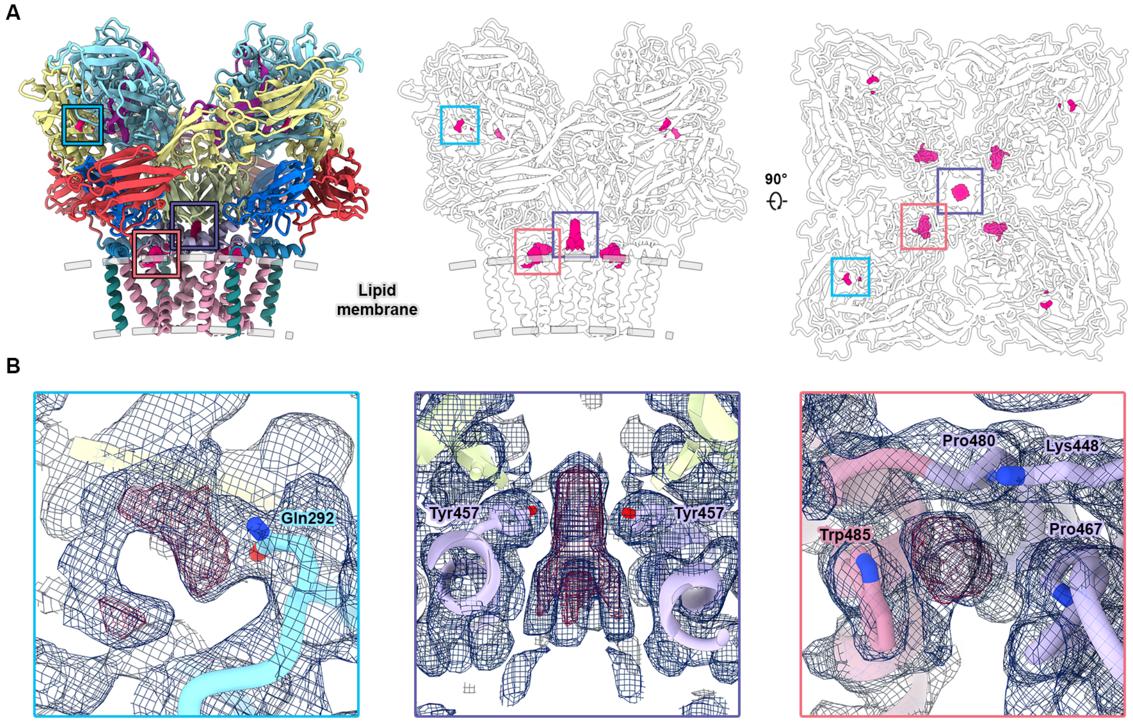


**Figure S4. Unidentified densities within the ANDV Gn–Gc tetramer map.**
(**A**) Structural model of the ANDV Gn–Gc tetramer overlaid with unidentified densities. Left: ribbon model colored by domain (color scheme as in Fig. 3). Right: outline-only representation of the model, demonstrating the spatial relationship between the unidentified densities (magenta surfaces) and the tetramer architecture. (**B**) Close-up views of the unidentified densities (pink mesh) embedded within the cryo-EM map (dark blue mesh). Sidechains of adjacent residues are shown as sticks to illustrate potential interactions.

### Figure S5

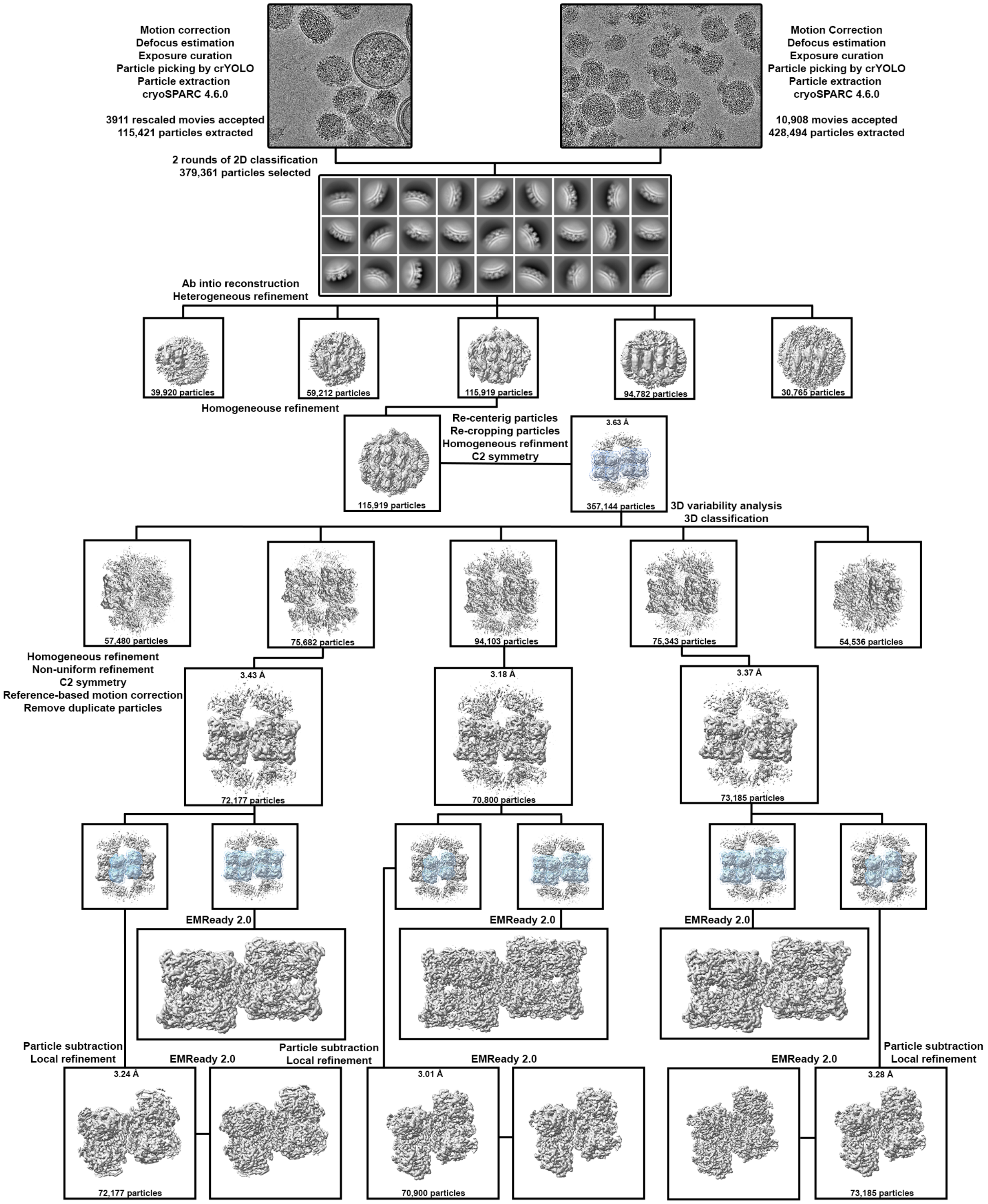


**Fig. S5. Cryo-EM data processing workflow for the ANDV Gn–Gc dimer of tetramers.**
Flowchart illustrating the cryo-EM data processing steps used to obtain the structure of the ANDV Gn–Gc dimer of tetramers. Particle counts at each stage are indicated, along with the corresponding resolutions of the 3D reconstructions. Masks used for recentering and focused 3D classification are represented as blue transparent surfaces.

#### Figure S6

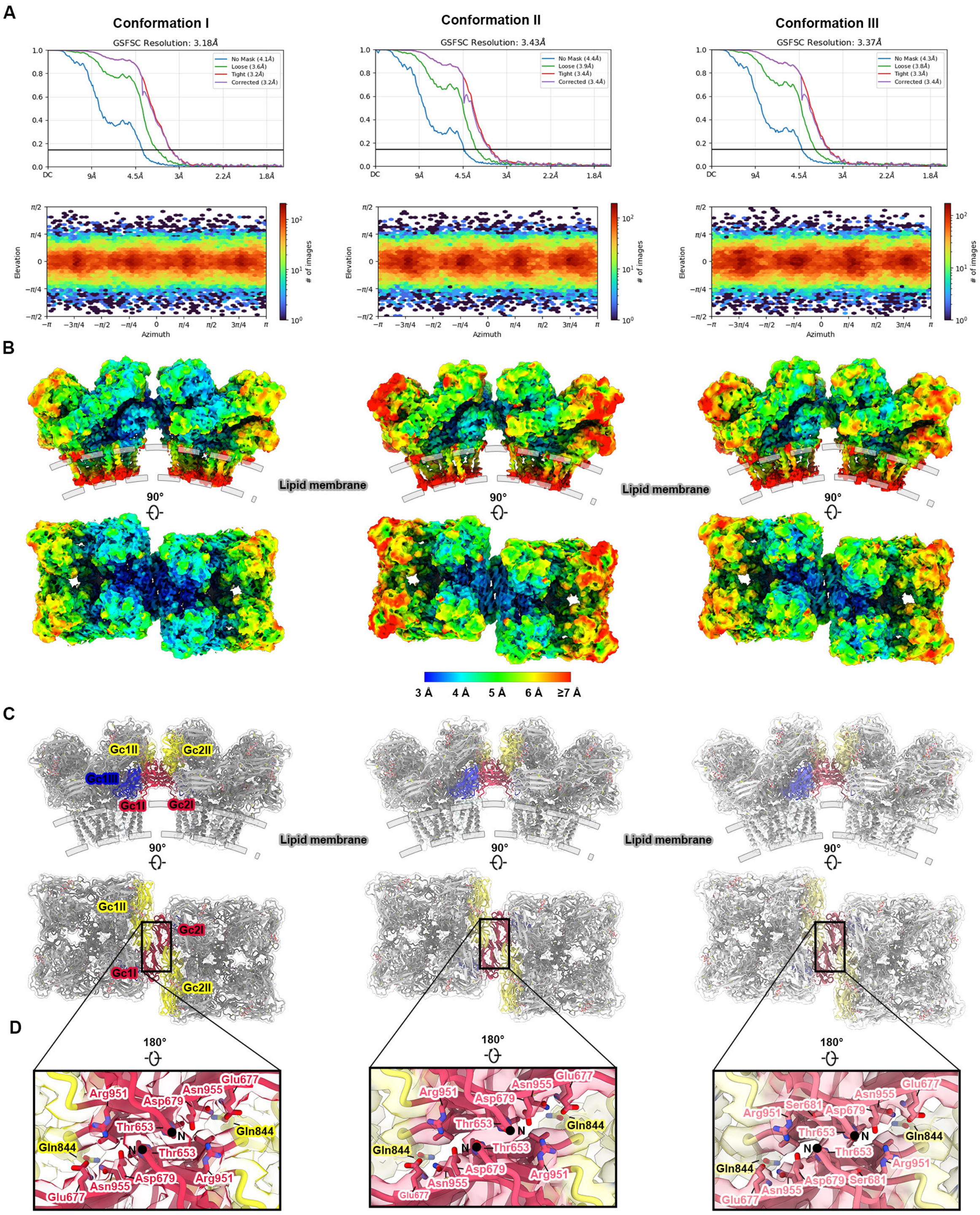


**Figure S6. Cryo-EM map validation and model fitting for the ANDV Gn–Gc dimer of tetramers.**
(**A**) Global gold-standard Fourier shell correlation (GSFSC) curves and particle orientation distribution plots for the three conformational states of the ANDV Gn–Gc dimer of tetramers (conformations I, II, and III). (**B**) Cryo-EM maps of the dimer of tetramers colored by local resolution. Transmembrane domains are positioned relative to a lipid bilayer, indicated by gray dashed curves. (**C**) Map-to-model fits for the three dimer conformations, highlighting the orientations of Gc homodimers formed between neighboring tetramers. (**D**) Detailed view of the tetramer–tetramer interface, showing the model fit within the EM map and the side chains of key interacting residues.

#### Figure S7

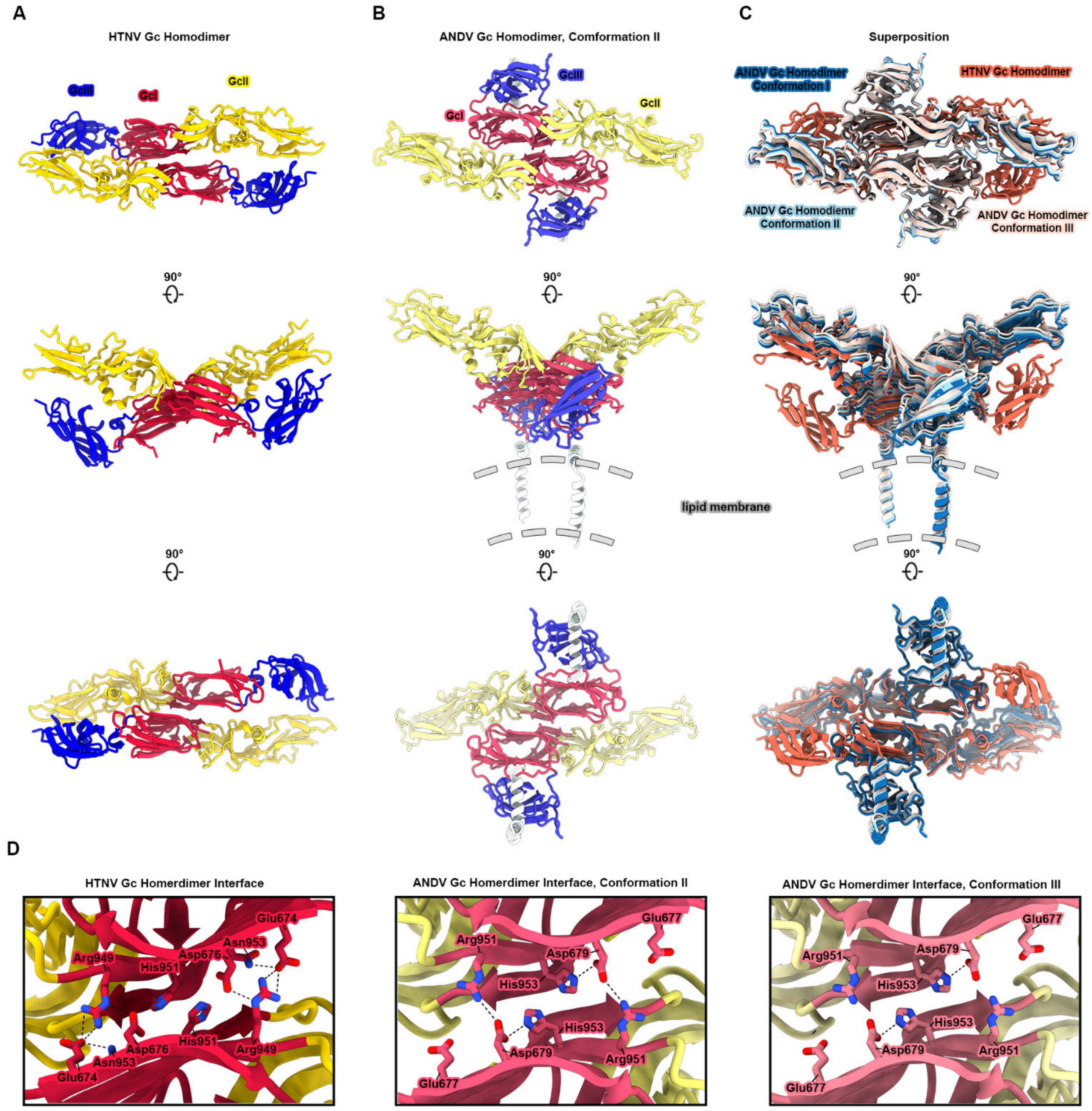


**Fig. S7. Comparison of the ANDV Gc homodimers with the Hantaan virus Gc crystallographic homodimer.**
(**A**) Different views of the Hantaan virus (HTNV) Gc crystallographic homodimer colored by domain. GcI, Gc domain I; GcII, Gc domain II; GcIII, Gc domain III. (**B**) Corresponding views of the ANDV Gc homodimer in conformation II, positioned relative to a lipid bilayer represented by gray dashed curves. (**C**) Structural superposition of ANDV Gc homodimers with the HTNV crystallographic dimer, aligned on Gc domain I. A slight downward rotation of one HTNV Gc protomer is observed, along with an extended conformation of HTNV Gc domain III. (**D**) Interface analysis highlights the absence of several hydrogen bond networks in the ANDV Gc homodimer that are present in the HTNV structure.

#### Figure S8

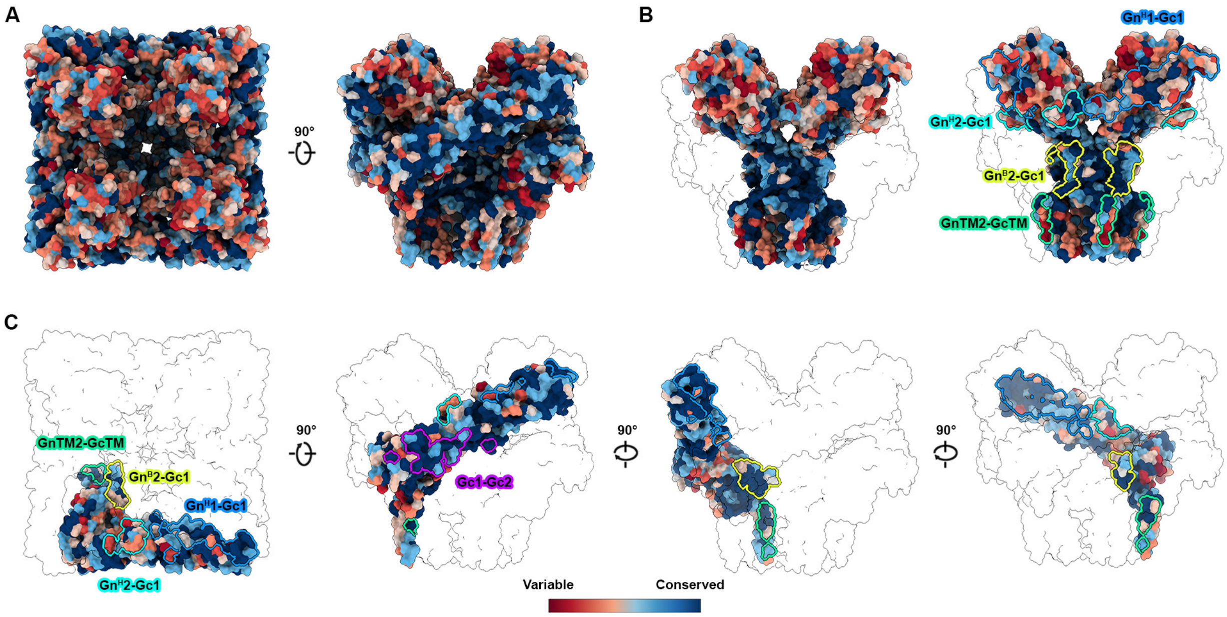


**Fig. S8. Conservation of hantavirus glycoprotein surfaces in the Gn–Gc tetramer**

Surface representations of the ANDV Gn–Gc tetramer (**A**), Gn tetramer (**B)**, and a single Gc protomer (**C**), colored by sequence conservation scores across representative hantaviruses (red: variable; blue: conserved). In panels B and C, other protomers are shown as surface outlines for context. A multiple sequence alignment was performed using 10 M-segment amino acid sequences with the highest hit scores for each of Hantaan virus (HTNV), Seoul virus (SEOV), Sin Nombre virus (SNV), ANDV, Puumala virus (PUUV), Dobrava-Belgrade virus (DOBV), and Tula virus (TULV). Key interfaces involved in tetramer stabilization and lattice organization are indicated.

#### Figure S9

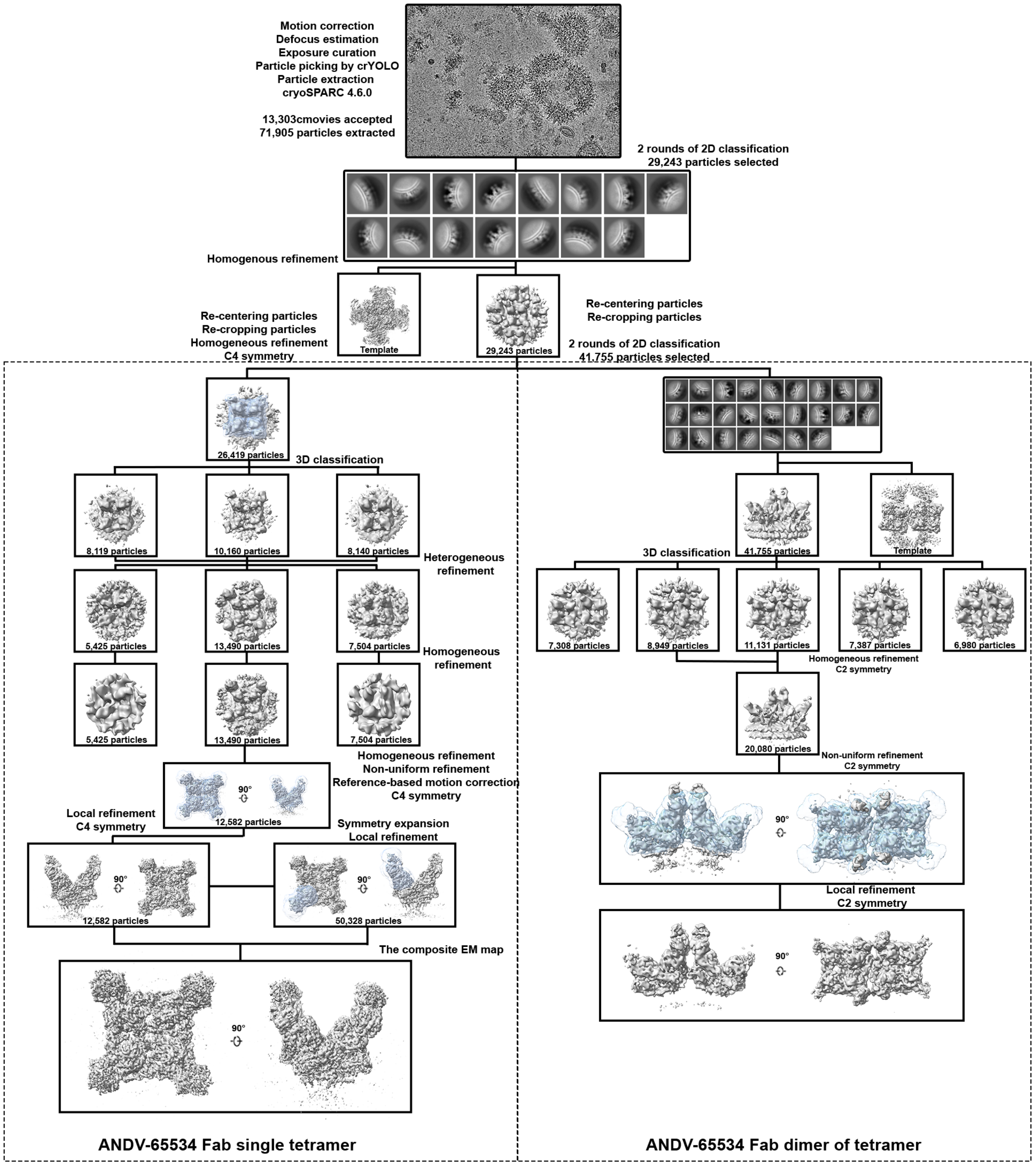


**Fig. S9. Cryo-EM data processing workflow for the ANDV-DM-eVLP complex with ADI-65534**

Flowchart illustrating the cryo-EM data processing steps used to obtain the structure of the ANDV-DM- eVLP complex with ADI-65534 Fab. Particle counts at each stage are indicated, along with the corresponding resolutions of the 3D reconstructions. Masks used for focused refinements are represented as blue transparent surfaces. Left: data processing for the ANDV Gn–Gc tetramer in complex with ADI-65534 Fab. Right: data processing for the ANDV Gn-Gc dimer of tetramers in complex with ADI-65534 Fab.

### Figure S10

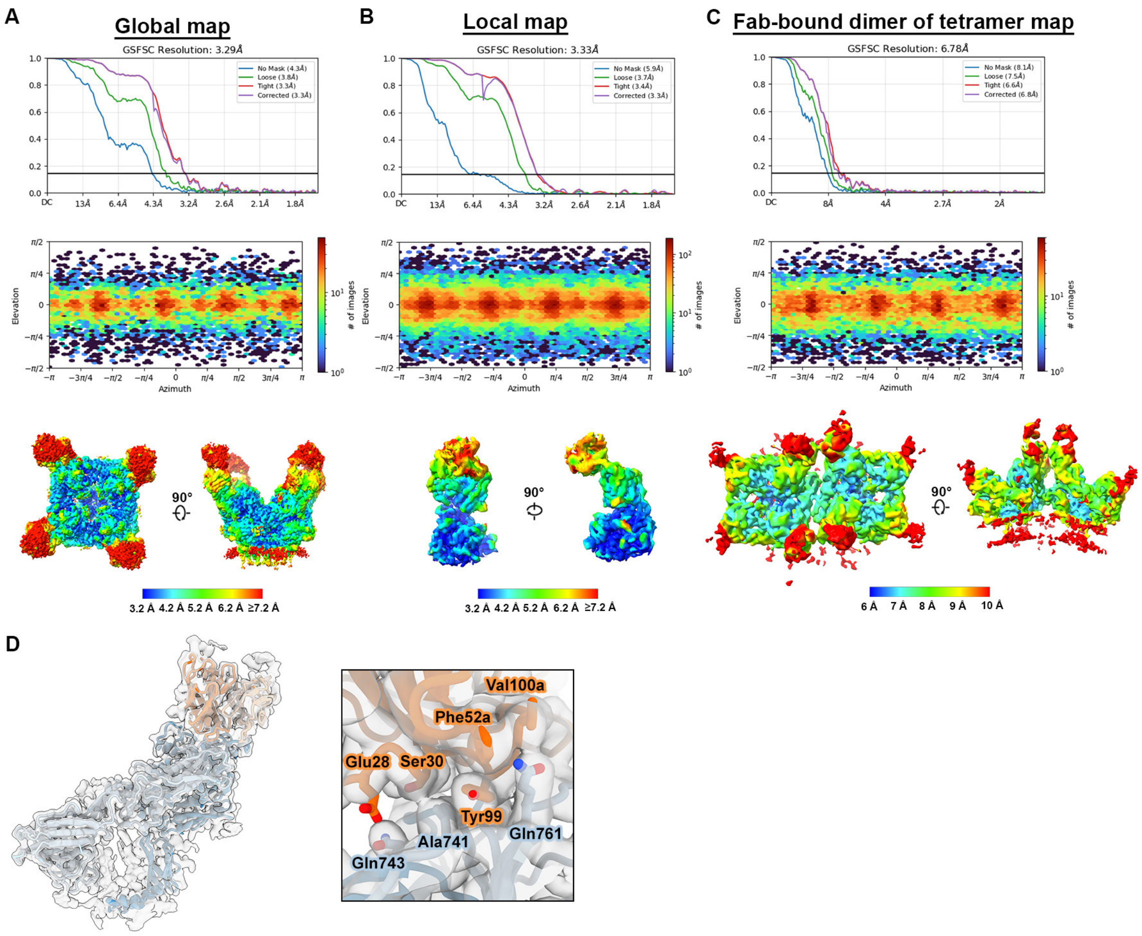


**Figure S10. Cryo-EM map validation and model fitting for the ADI-65534-bound ANDV Gn–Gc tetramer and dimer of tetramers.**

(A–C) Global gold-standard Fourier shell correlation (GSFSC) curves, particle orientation distribution plots, and cryo-EM maps colored by local resolution for: (**A**) the globally refined single tetramer map, (**B**) the locally refined map focused on the Fab interface, and (**C**) the dimer of tetramers map. (**D**) Detailed view of the Gn–Gc-Fab interface, showing the model fit within the cryo-EM map and the sidechains of key interacting residues.

#### Figure S11

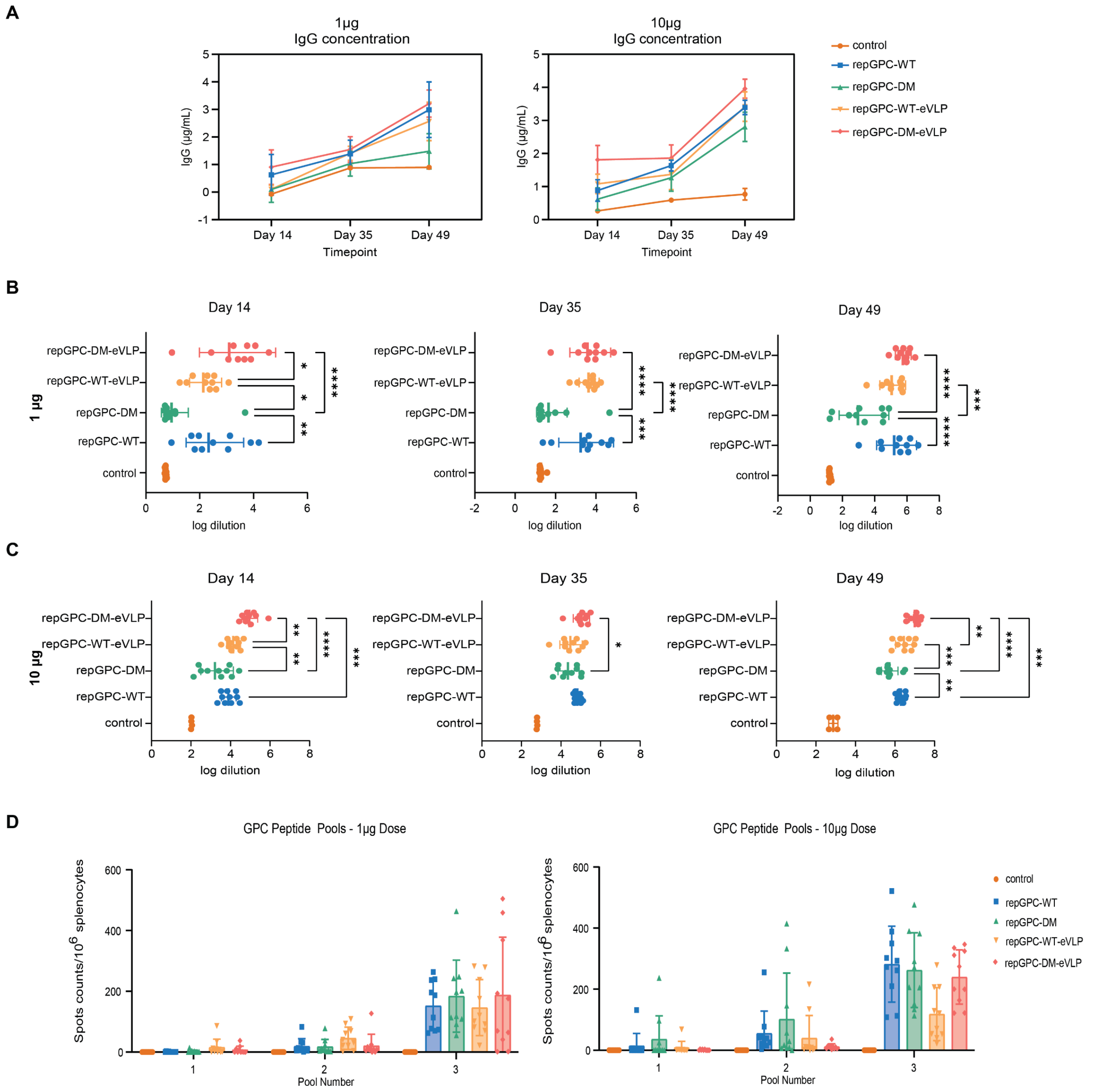


**Fig. S11. Humoral and cellular immune response elicited by ANDV-GPC constructs**

(**A**) Time course of serum IgG levels against ANDV Gn–Gc following immunization with 1 μg (left) or 10 μg (right) of repRNA/LIONTM ANDV-GPC constructs. (**B–C**) Endpoint titers of Gn–Gc-specific antibodies on Days 14, 35, and 49 post-immunization with 1 μg (B) or 10 μg (C) of repRNA/LIONTM ANDV-GPC constructs. Titers were defined as the reciprocal serum dilution yielding an absorbance ≥2 standard deviations above the background. (**D**) IFN-γ ELISpot responses in splenocytes from mice immunized with 1 μg (left) or 10 μg (right) of repRNA/LIONTM ANDV-GPC constructs and stimulated with pooled ANDV-GPC peptides. Spot counts are shown per $10^6$ splenocytes after background subtraction. Data in all panels are presented as mean or geometric mean with standard deviation. Statistical analysis was conducted using one-way ANOVA with Tukey's multiple comparisons test.
